# Loss-of-function of *OTUD7A* in the schizophrenia-associated 15q13.3 deletion impairs synapse development and function in human neurons

**DOI:** 10.1101/2022.01.06.473910

**Authors:** Alena Kozlova, Siwei Zhang, Alex V. Kotlar, Brendan Jamison, Hanwen Zhang, Serena Shi, Marc P. Forrest, John McDaid, David J. Cutler, Michael P. Epstein, Michael E. Zwick, Zhiping P. Pang, Alan R. Sanders, Stephen T. Warren, Pablo V. Gejman, Jennifer G. Mulle, Jubao Duan

## Abstract

Identifying causative gene(s) within disease-associated large genomic regions of copy number variants (CNVs) is challenging. Here, by targeted sequencing of genes within schizophrenia (SZ)-associated CNVs in 1,779 SZ cases and 1,418 controls, we identified three rare putative loss-of-function (LoF) mutations in OTU deubiquitinase 7A (OTUD7A) within the 15q13.3 deletion in cases, but none in controls. To tie OTUD7A LoF with any SZ-relevant cellular phenotypes, we modeled the OTUD7A LoF mutation, rs757148409, in human induced pluripotent stem cell (hiPSC)-derived induced excitatory neurons (iNs) by CRISPR/Cas9 engineering. The mutant iNs showed a ∼50% decrease in OTUD7A expression without undergoing nonsense-mediated mRNA decay. The mutant iNs also exhibited marked reduction of dendritic complexity, density of synaptic proteins GluA1 and PSD-95, and neuronal network activity. Congruent with the neuronal phenotypes in mutant iNs, our transcriptomic analysis showed that the set of OTUD7A LoF-downregulated genes was enriched for those relating to synapse development and function, and was associated with SZ and other neuropsychiatric disorders. These results suggest that OTUD7A LoF impairs synapse development and neuronal function in human neurons, providing mechanistic insight into the possible role of OTUD7A in driving neuropsychiatric phenotypes associated with the 15q13.3 deletion.

## INTRODUCTION

Genome-wide association studies (GWAS) of schizophrenia (SZ) and other neuropsychiatric disorders have identified hundreds of risk loci with common genetic risk variants (Consortium, 2011; Consortium et al., 2020; Consortium., 2014; Mullins et al., 2021; Purcell et al., 2009; Ripke and Consortium., 2013; Shi et al., 2009; Stefansson et al., 2009). However, such common disease risk variants often have small effect sizes and lie in noncoding regions of the genome, hindering the mechanistic understanding of disease pathophysiology. These genome-wide studies also revealed another side of the risk spectrum for neuropsychiatric and neurodevelopmental disorders (NDDs): rare copy number variants (CNVs) that are strongly associated with SZ and have much larger effect sizes (odds ratios, OR of 2–70) than common SNPs (OR <1.2) (Bassett et al., 2010; Levinson et al., 2011; Marshall et al., 2017; Szatkiewicz et al., 2014). CNVs reproducibly associated with SZ include deletions at 1q21.1, 2p16.3 (*NRXN1*), 3q29, 15q13.3, distal 16p11.2, and duplications at 7q11.23 and proximal 16p11.2 (Bassett et al., 2010; Levinson et al., 2011; Marshall et al., 2017; Szatkiewicz et al., 2014). The large effect sizes of these variants make them an ideal model for understanding disease biology and interpreting the disease relevance of cellular phenotypes.

Rare SZ-associated CNVs are usually long (>100 kb) and span multiple genes (Bassett et al., 2010; Levinson et al., 2011; Marshall et al., 2017; Szatkiewicz et al., 2014). Furthermore, CNV- carriers often manifest heterogeneous clinical phenotypes. It has been a challenge to identify which gene(s) within a CNV region are likely the driver(s) for disease-relevant phenotypes. The challenge is amplified by possible effects of a CNV on local or distal chromatin architecture and, subsequently, the expression of genes outside the CNV region (Franke et al., 2016; Redin et al., 2017). As a result, the disease-relevant cellular phenotypes and the driver gene(s) for most CNVs remain elusive.

Individuals who harbor the SZ-associated 15q13.3 deletion (Hasselmo, 2006; Levin, 2012; Soler-Alfonso et al., 2014) are susceptible to a wide range of NDDs including SZ and autism spectrum disorder (Freedman et al., 2001; Gillentine and Schaaf, 2015; Helbig et al., 2009; Pagnamenta et al., 2009; Sharp et al., 2008; Shinawi et al., 2009; Stefansson et al., 2008; Ziats et al., 2016). Initially, the *cholinergic receptor nicotinic alpha 7 subunit* (*CHRNA7*) in the 15q13.3 deletion was favored as a likely driver gene because it encodes a neuronal ligand-gated ion channel. This superfamily of ion channels mediate fast signal transmission at synapses, synaptic plasticity, and transduction (Hasselmo, 2006; Levin, 2012; Soler-Alfonso et al., 2014). However, mice deficient in *Chrna7* do not recapitulate the behavioral and cognitive phenotypes observed in humans (Yin et al., 2017). Instead, the knockout mice for the ortholog of another gene within the CNV region, *Otud7a*, recapitulate cardinal phenotypes associated with the 15q13.3 deletion syndrome, such as developmental delay, vocal impairment, seizure-related abnormalities, motor deficits, hypotonia, and acoustic startle deficit (Yin et al., 2018). *Otud7a* encodes a putative deubiquitinating enzyme that localizes to spine compartments, and *Otud7a*- null mice showed decreased dendritic spine density as well as reduced frequency of miniature excitatory postsynaptic currents (Yin et al., 2018). In addition, a 5-year-old female with developmental delay was found to have a genetic deletion spanning *OTUD7A*, but not *CHRNA7* (Uddin et al., 2018), further suggesting that haploinsufficiency of *OTUD7A* likely drives some critical phenotypes of the 15q13.3 deletion syndrome.

It is currently unknown whether haploinsufficiency of *OTUD7A* in humans may result in cellular phenotypes relevant to SZ or other NDDs. Here, we sequenced 241 genes within SZ-associated CNVs in a cohort of SZ cases and unaffected controls to find rare protein-coding variants. We identified three putative *OTUD7A* loss-of-function (LoF) mutations that were only present in SZ cases. Human induced pluripotent stem cell (hiPSC)-derived neurons, combined with CRISPR/Cas9 editing, have been used to create isogenic cellular models for dissecting the functional impacts of both common GWAS risk variants (Forrest et al., 2017; Schrode et al., 2019; Zhang et al., 2020) and rare genetic variants associated with SZ and other neuropsychiatric disorders (Duan et al., 2019; Flaherty et al., 2019; Pak et al., 2021a; Wen et al., 2016; Zhang et al., 2021). Thus, we further modeled the *OTUD7A* LoF mutation by CRISPR/Cas9 editing of human iPSC-derived excitatory neurons (Zhang et al., 2013). We found that *OTUD7A* LoF reduced protein expression, complexity of neuronal arbors, and impaired development of glutamatergic synapses and neural network activity. An integrative analysis of isogenic neural transcriptomes and neuropsychiatric GWAS further showed that *OTUD7A* LoF downregulated genes associated with synapse development and function, as well as with SZ and other neuropsychiatric disorders. These data advance *OTUD7A* as a driver of neurodevelopmental and psychiatric phenotypes at the 15q13.3 deletion.

## RESULTS

### Targeted resequencing of CNV genes identifies *OTUD7A* rare LoF mutations only in SZ patients

To identify disease-relevant protein-coding rare variants in genes within 6 SZ-associated CNV intervals (1q21 del, 3q29 del, 7q11.23 dup, 15q13.3 del, 16p11.2 dup, 22q11.2 del) (Bassett et al., 2010; Levinson et al., 2011; Marshall et al., 2017; Szatkiewicz et al., 2014), we performed a targeted exon resequencing of all CNV genes in 1,779 SZ cases and 1,418 controls from the Molecular Genetics of SZ (MGS) cohort (Sanders et al., 2010; Shi et al., 2009). Constrained by the relatively small sample size, no individual variants reached significant association with SZ. Furthermore, rare deleterious variants grouped by their corresponding CNV interval also failed to show significant association (data not shown). Because gene LoF as a result of protein stop codon gain (i.e., stop-gain, nonsense mutations) and frameshift insertion/deletion mutations could mimic the deleterious effects of a CNV deletion, we further examined whether any CNV genes had rare (MAF < 0.005) LoF mutations associated with SZ. Although no significant associations were identified for individual rare LoF mutations (Table S1) or LoF variants grouped by individual genes (Table S2), *OTUD7A* in the 15q13.3 deletion region was ranked as the strongest candidate among all LoF-intolerant genes (probability of LoF intolerance score, pLi >= 0.9) (Figure 1A, Table S2).

**Fig. 1:**
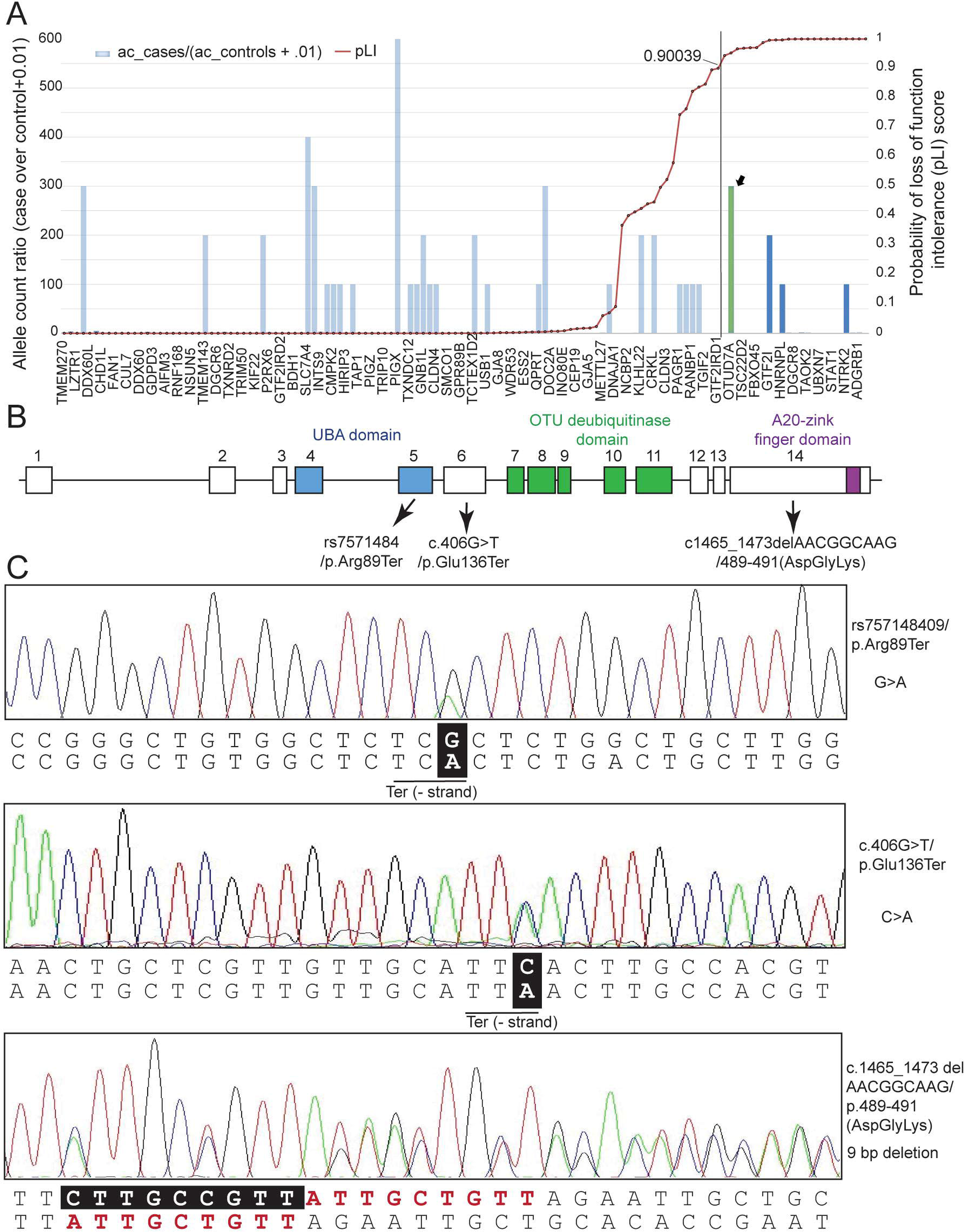
Evidence for the association of *OTUD7A* with SZ. (A) The case/control ratio of LoF (stop gain and frameshift) alleles among samples with complete genotypes. *OTUD7A* had the strongest evidence of a LoF mutational bias among LoF-intolerant genes (pLi ≥ 0.9), with 3 alleles identified in cases and none in controls. **(B)** Structure of *OTUD7A*. Exons of *OTUD7A* are boxed, whereas introns are represented by lines in between exons. All exons and introns are plotted in scale within their category. Three domains have been identified in *OTUD7A*: the UBA domain (blue), the OTU deubiquitinase domain (green), and an A20 zinc finger domain (purple). Three mutations identified in SZ patients in exons 5, 6 and 14 are indicated. **(C)** Sanger sequencing confirmation of the putative three LoF mutations identified in SZ patients. Two of these mutations are SNPs (top and middle sequences) which result in premature stop codons (rs757148409/p.Arg89Ter and c.406G>T/p.Glu136Ter, respectively), while the third mutation (bottom sequence) is a 9 bp in-frame deletion (c.1465_1473delAACGGCAAG/p.489- 491(AspGlyLys)).

We found three putative LoF mutations or protein-truncating variants (PTVs) in exons 5, 6, and 14 of *OTUD7A* in 3 SZ cases, but none in controls (Figure 1B). *OTUD7A* encodes a protein with three domains: a ubiquitin-associated (UBA) domain, an ovarian tumor (OTU) deubiquitinase domain, and an A20-like zinc finger domain (Figure 1B). Two of these mutations (in exons 5 and 6) are SNPs, which result in premature stop codons (rs757148409/p.Arg89Ter and c.406G>T/p.Glu136Ter, respectively) at the UBA domain and were confirmed by Sanger sequencing (Figure 1C). While the third mutation was initially called as a 5-bp frameshift deletion (in exon 14; c.1465_1469delAACGG/p.758Ter) at the A20-like zinc finger domain, it was shown by Sanger sequencing to be a 9-bp in-frame deletion (c.1465_1473delAACGGCAAG/p.489-491(AspGlyLys)) (Figure 1C). All three carriers of these rare mutations had chronic SZ (>30 years illness) with typical young adulthood onset (19-22 years old in these subjects) (Table S3). One subject had an unspecified learning disorder and only completed the ninth grade; the other two subjects completed the tenth grade (and later got a GED) or the twelfth grade (and graduated high school). By further examining the SZ Exome Sequencing Meta-analysis (SCHEMA) database with 24,248 SZ cases and 97,322 population controls (Singh et al., 2020), we found only rs757148409/p.Arg89Ter, at a SCHEMA allele frequency of 0.000013 (Table S4). We did not find significant association of SZ with *OTUD7A* LoF mutations in SCHEMA; however, combining the LoF mutations in our resequencing study and SCHEMA yielded a non-significant enrichment of rare *OTUD7A* LoF mutations in SZ cases (Fold enrichment = 2.7, Fisher’s Exact Test *p*-value = 0.12) (Tables S3 and S4). Given the high pLI score (0.95) for *OTUD7A* (Karczewski et al., 2020), it is conceivable that the rare *OTUD7A* LoF mutations identified in our SZ patients are likely to have deleterious effects. Also, in consideration of a study of *Otud7a*-null mice that suggested that haploinsufficiency of *OTUD7A* may drive phenotypes of the 15q13.3 deletion syndrome (Yin et al., 2018), we prioritized *OTUD7A* among all the sequenced CNV genes for further modeling the impact of LoF in human neurons.

### *OTUD7A* LoF mutation leads to a reduced protein expression in human neurons

We next modelled the LoF of *OTUD7A* in hiPSC-derived induced excitatory neurons (iNs), aiming to test whether *OTUD7A* LoF was associated with SZ-relevant neurodevelopmental phenotypes in human neurons. We first performed CRISPR/Cas9 editing (Ran et al., 2013) to introduce the risk allele of an LoF mutation (rs757148409) into iPSC lines derived from two control (i.e., non-SZ) donors. We selected this mutation for functional study because it was in exon 5 and the resultant truncating protein would be the shortest among the three identified mutations in SZ patients. For each line, we generated three isogenic pairs of hiPSC clones that carried a heterozygous G to A mutation at rs757148409, resulting a stop-codon-gain (Figure 2A- B). The presence of the heterozygous on-target editing and the absence of the predicted top-ranking off-target editing were confirmed by Sanger sequencing (Figures 2B and S1A). The isogenic pairs of CRISP/Cas9-edited hiPSC lines were also characterized for pluripotency by immunofluorescence staining (Figure S1B), and the absence of chromosomal abnormality was verified by RNA-seq-based e-karyotyping (Zhang et al., 2020) (Figure S1C).

**Fig. 2:**
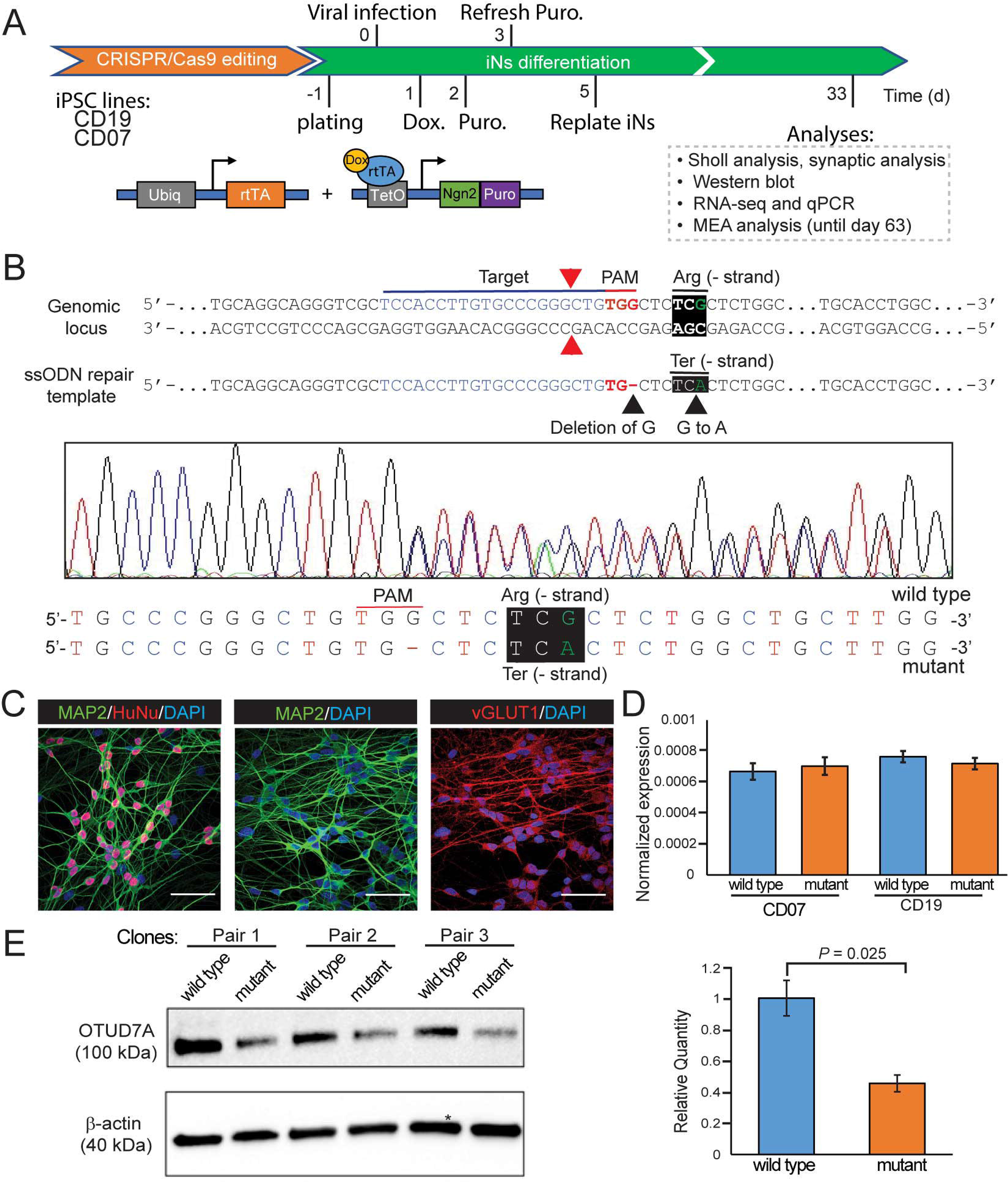
***OTUD7A* LoF mutation leads to a reduced protein expression in human neurons. (A)** Experimental design for generating and analyzing NGN2-induced neurons from hiPSCs. Ubiq, ubiquitin; rtTA, reverse tetracycline-controlled transactivator; Puro, puromycin; Dox, Doxycycline. **(B)** Schematics of CRISPR/Cas9 editing at SNP rs757148409. The Cas9 nuclease is targeted to genomic DNA by a sgRNA consisting of 20-nt guide sequence. The guide sequence pairs with the DNA target directly upstream of a requisite 5’-NGG motif (PAM: red). Cas9 mediated a DSB ∼3bp upstream of the PAM (red triangle). G was deleted from the PAM sequence (black triangle) in ssODN repair oligo and G was replaced with A at SNP rs757148409 (black triangle). Two control homozygous GG (Arg) cell lines were used to generate heterozygous GA (Ter) at the SNP site introducing stop-codon-gain. For each line, three isogenic pairs of hiPSC clones were generated. Shown is a + strand sequence, but mRNA is transcribed from the – strand. SNP editing was confirmed by Sanger sequencing in sub- cloned CRISPR-edited hiPSC clones. **(C)** Representative immunofluorescence images of isogenic iNs (MAP2+/VGLUT1+). Scale bar: 50 µm. **(D)** Bar graph of qPCR showing that the mRNA level of *OTUD7A* was not reduced in the mutant iNs compared to wild type iNs in both CD07 and CD19 lines. Two biological replicates were included for each experimental group, and three technical replicates were included in qPCR. Expression was normalized to GAPDH. Error bars are SEM. **(E)** Western blotting with OTUD7A antibodies and quantification showing that the expression of OTUD7A protein (100 kDa) was significantly reduced (*p* = 0.025) by ∼50% in the mutant CD19 iNs. Student’s t-test was used to quantify statistical significance. Error bars are SEM.

Based on the predominant brain expression of *OTUD7A* in the GTEx database (Consortium, 2013), and its more robust expression in glutamatergic (excitatory) neurons as observed in the Allen Brain Map (Yao et al., 2021) and in our hiPSC-derived neurons (Zhang et al., 2020) (Figure S2A-C), we differentiated the isogenic pairs of CRISPR/Cas9-edited hiPSC lines into the iNs by infecting the hiPSCs with a lentivirus cocktail containing NGN2 and rtTA viruses (Zhang et al., 2013) (Figure 2A). We obtained iNs (PSD-95+, VGLUT1+, MAP2+, and HuNu+) (Figure 2C) of relatively high purity (∼96%; Figure S3A,B). We also confirmed the functionality of these iNs by examining their action potentials (Figure S3C).

With these isogenic iNs, we first examined whether the CRISPR-engineered premature stop-gain mutation rs757148409/p.Arg89Ter in *OTUD7A*, like most pathogenic premature stop-gain mutations in ClinVar, caused complete LoF of the mutant allele through triggering nonsense-mediated mRNA decay (NMD) (Supek et al., 2021). Our quantitative PCR (qPCR) result showed that the mRNA level of *OTUD7A* was not reduced in the mutant iNs compared to wild type iNs (Figure 2D). This suggested that the mutant allele of rs757148409 of *OTUD7A* gene did not cause NMD, but rather likely resulted in truncated proteins/peptides. Our Western blotting with OTUD7A antibody further showed that the expression of the wild type OTUD7A protein was reduced by about 50% in the mutant iNs (Figure 2E). We did not observe on the Western blot a much smaller truncated protein/peptide (p.Arg89Ter; ∼10kD) presumably produced from the mutant allele; however, given its short length and the lack of the OTU deubiquitinase domain, it is conceivable that the truncated protein/peptide was likely not functional. Nonetheless, the heterozygous rs757148409/p.Arg89Ter of *OTUD7A* identified in our SZ patient likely resulted in LoF of *OTUD7A*, which would be reminiscent of the functional effect of the SZ-associated 15q13.3 deletion if *OTUD7A* is a “driver” gene within the CNV as previously suggested by studying an animal model (Yin et al., 2018).

### *OTUD7A* LoF mutation impairs neurodevelopmental phenotypes in human neurons

To evaluate the effects of the *OTUD7A* LoF mutation on neurodevelopmental phenotypes in human neurons (Figure 2A), we first examined the alteration of dendritic complexity in isogenic iNs carrying the heterozygous rs757148409/p.Arg89Ter. We sparsely transfected iNs on day 28 with green fluorescent protein (GFP)-expressing plasmid DNAs to visualize dendritic morphology and measured dendritic arbors using Sholl analysis. We found significant differences in dendritic morphology between the mutant and wild type CD07 (Figure 3A) and CD19 (Figure 3B) iNs lines. The mutant iNs exhibited fewer branches than wild type iNs.

**Fig 3.**
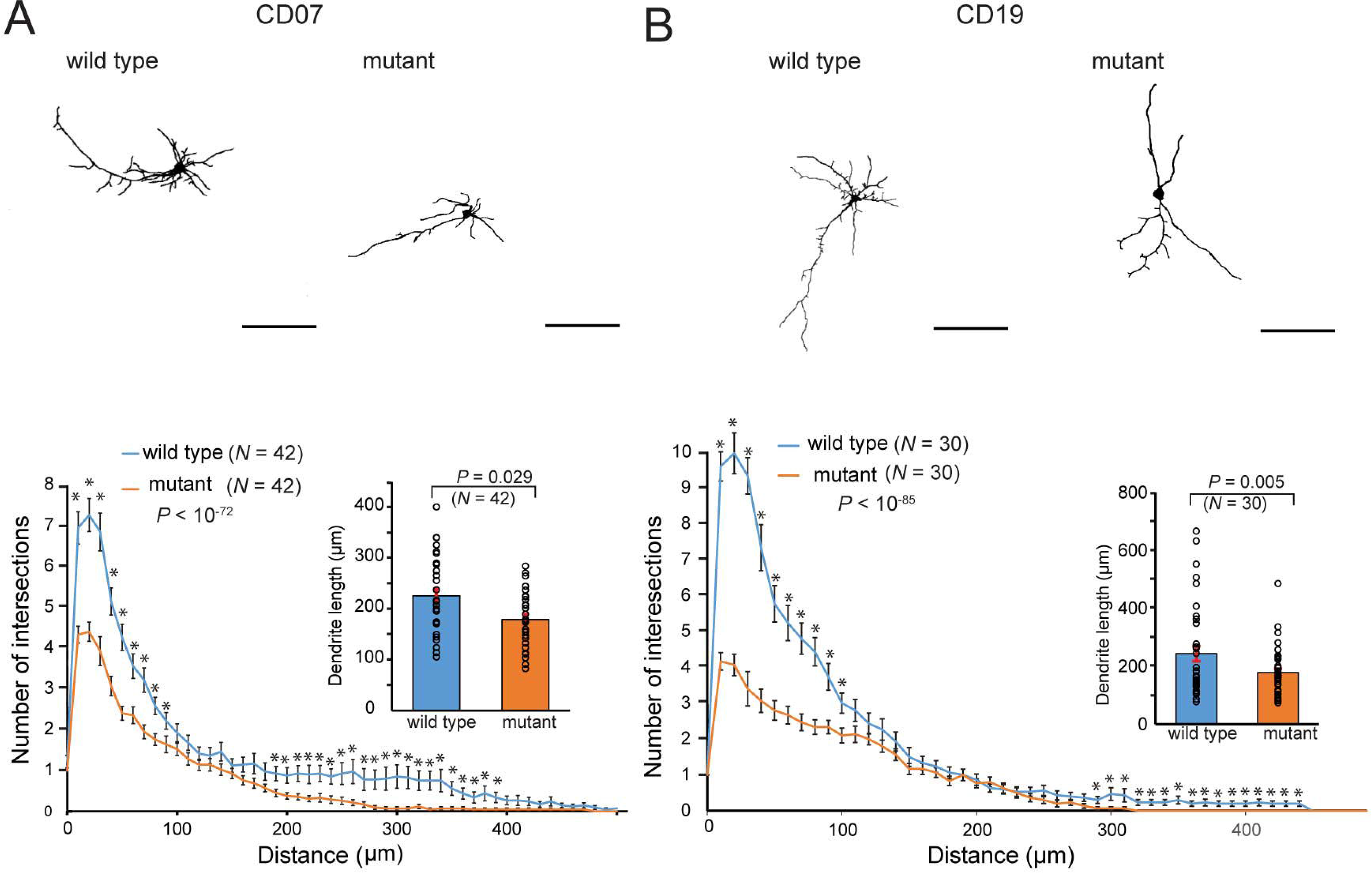
*OTUD7A* LoF mutation alters dendritic complexity. Representative binary traces of GFP transfected iN-d34, sholl analysis and dendrite length for *OTUD7A* mutant and wild type CD07 (**A**) and CD19 (**B**) lines. Scale bar: 100 µm. Sholl analysis of total dendritic branching shows a significant ∼50% reduction in branching near soma and the distal end in mutant CD07 (n=42, *p* < 10^-72^) and CD19 (n=30, *p* < 10^-85^) iNs compared with the wild type lines. Two-way repeated-measures ANOVA was used to quantify overall statistical significance between mutant and wild type lines. Student’s t-test was used to quantify significant difference between mutant and wild type lines at each distance point, \**p* < 0.05. *OTUD7A* mutant iNs show ∼30% decrease in total dendrite length in both CD07 (n=42, *p* <10^-2^) and CD19 (n=30, *p* < 10^-3^) lines when compared with wild type iNs. Circles represent individual data points for each genotype. Error bars are SEM. Student’s t-test was used to quantify statistical significance.

Significant differences of dendritic branching were observed near the soma (<100 μm) and in the distal end (>200-300 μm). There was ∼50% reduction in dendritic complexity and ∼30% decrease in total dendrite length in the mutant iNs compared with the wild type lines. These results suggested that *OTUD7A* LoF mutation was associated with decreased neuronal complexity and had effects on the dendritic development of human iNs, Glutamatergic synapses are the main excitatory synapses in the brain and have been implicated in the pathophysiology of psychiatric disorders (Volk et al., 2015). To determine whether *OTUD7A* LoF mutation can affect glutamatergic synapse development, we immunostained the GFP-transfected iNs with PSD-95, a postsynaptic scaffolding protein, and GluA1, a subunit of α-amino-3-hydroxy-5-methyl-4-isoxazolepropionic acid (AMPA)-type glutamate receptor, both of which accumulate in mature glutamatergic synapses (Meyer et al., 2014). We found that both markers formed characteristic puncta on dendrites and in dendritic protrusions of human iNs (Figure 4A,B). We measured density and area of GluA1 puncta, PSD-95 puncta, or both proteins co-localized (GluA1+/PSD-95+ puncta) along dendritic branches. For both lines, we found that the mutant iNs showed a marked decrease of both GluA1 and PSD-95 puncta (Figure 4C,D), and a concomitant reduction in the density and area of overlapping of GluA1+/PSD-95+ puncta (Figure 4E), compared with wild type lines.

**Fig 4.**
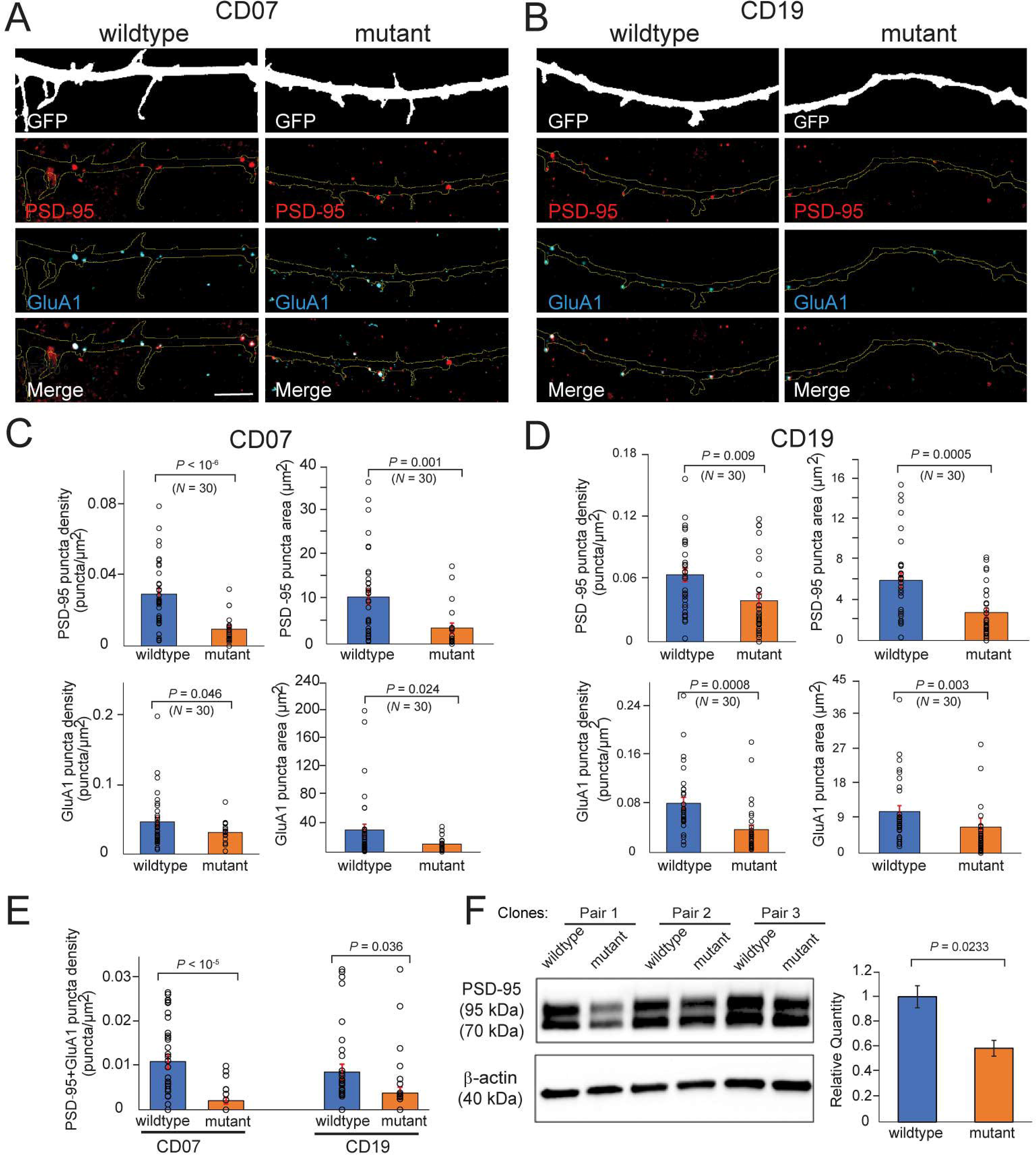
*OTUD7A* LoF mutation affects the maturity of dendritic protrusions. Composite confocal images of dendrites from GFP-transfected induced neurons (iN-d34) immunostained for PSD-95 and GluA1 in mutant and wild type iNs CD07 **(A)** and CD19 **(B)** lines. Both markers formed characteristic puncta on dendrites and in dendritic protrusions of neurons. PSD-95 puncta density and area is significantly decreased in CD07 **(C, top panel)** (n=30, *p* < 10^-6^ and n=30, *p* = 0.001 respectively) and CD19 **(D, top panel)** (n=30, *p* = 0.009 and n=30, *p* < 10^-3^ respectively) mutant iNs compared with the wild type lines. vGluA1 puncta density and area is significantly decreased in CD07 **(C, bottom panel)** (n=30, *p* = 0.046 and n=30, *p* = 0.024 respectively) and CD19 **(D, bottom panel)** (n=30, *p* = 0.024 and n=30, *p* = 0.003 respectively) mutant iNs when compared with wild type iNs. **(E)** GluA1+/PSD-95+ puncta density was significantly reduced in both CD07 and CD19 mutant iNs compared to wild type lines. Circles represent individual data points for each genotype in B-D. **(F)** Western blotting with PSD-95 antibodies and quantification showing that the expression of PSD-95 protein (two bands of 95kDa and 70kDa) was significantly reduced (*p* = 0.233) by ∼50% in the mutant CD19 iNs. Student’s t-test was used to quantify statistical significance. Error bars are SEM.

Consistent with the decrease in PSD-95 puncta density and area in the mutant lines, our Western blotting with PSD-95 antibodies also showed that PSD-95 protein expression (both the 95 kDa and the 70 kDa bands; the lower molecular weight band representing a truncated form of PSD-95 lacking the N terminus (Cho et al., 1992; Kornau et al., 1995)) was decreased by approximately 50% in the mutant iNs (Figure 4F). Taken together, our results suggest that the rs757148409/p.Arg89Ter mutation identified in our SZ patient is a LoF mutation, reducing dendritic complexity and glutamatergic synapse development in human iNs.

### *OTUD7A* LoF mutation reduces neural network activity

We reasoned that the observed neurodevelopmental deficits in iNs carrying the *OTUD7A* LoF mutation may impair neuronal and synaptic function. To test this hypothesis, we measured the neuronal network activity at the population level by multi-electrode array (MEA) in mutant and wild type CD07 (Figure 5A) and CD19 (Figure 5B) iNs lines. For different clones of the two isogenic pairs of hiPSC lines, we re-plated iNs (co-cultured with glial cells) on MEA plates and recorded neuronal activity from day 33 to day 63 post neural induction (Figure 2A). We found that the neural firing frequency, number of bursts, and network burst frequency in iNs carrying the *OTUD7A* LoF mutation exhibited significant decreases compared to the wild type neurons (Figure 5A,B and S4A,B), which were consistent between neurons of both hiPSC lines. The iNs from the wild type hiPSC lines showed their highest firing frequency at about day 41 and network burst frequency at about day 44-48 (Figure S4A,B), when iNs also showed the most robust differences for these metrics between the wild type and the mutant neurons (Figure 5C,D). The consistent reduction of neuronal network activity in the mutant lines of two different donors or genetic backgrounds suggests a highly penetrant effect of the *OTUD7A* LoF mutation on neuronal activity.

**Fig 5.**
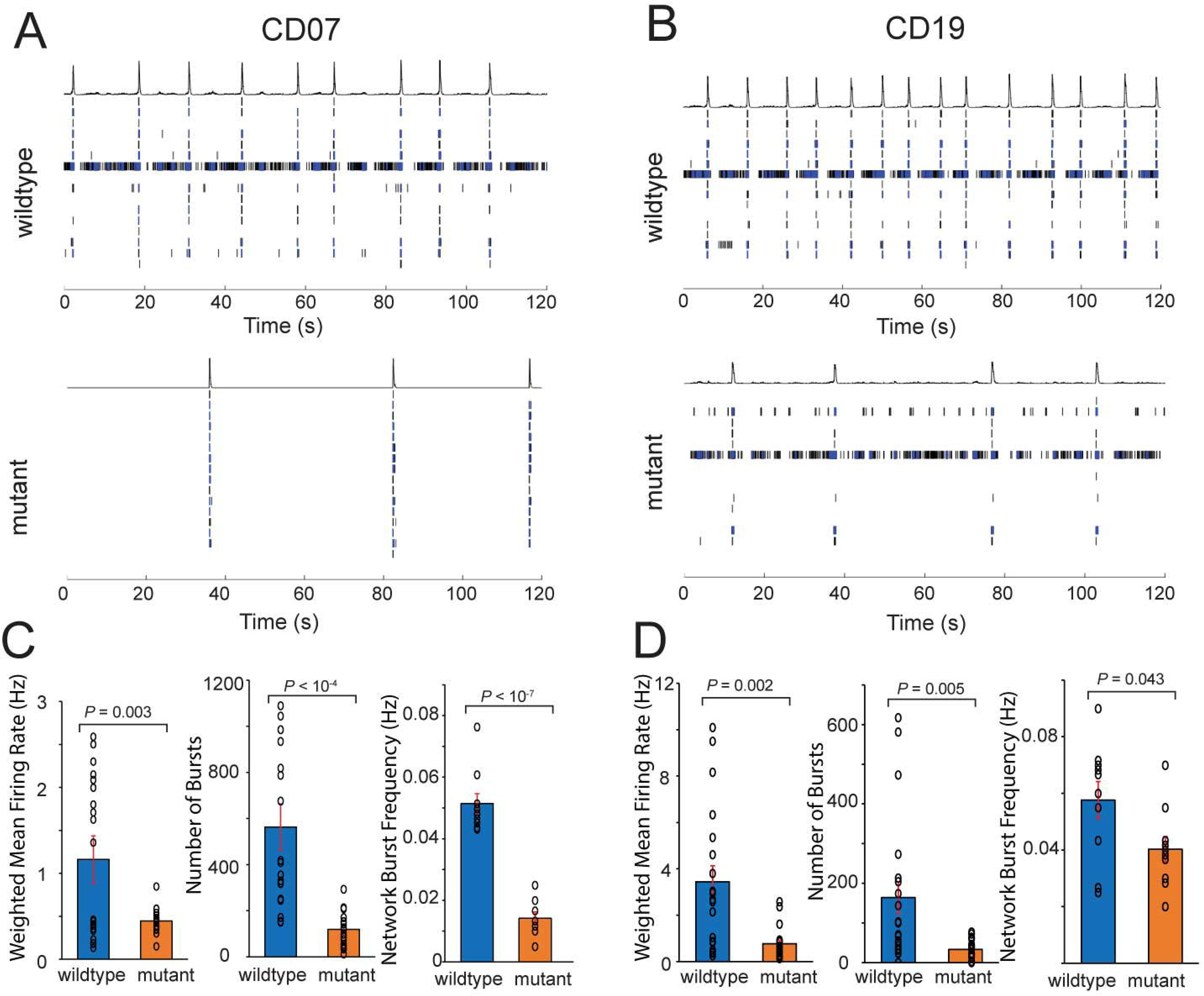
*OTUD7A* LoF mutation reduces neural network activity. Representative raster plot of electrophysiological activity exhibited by wild type and mutant iNs’ network at different time points in CD07 (**A**) and CD19 (**B**). Quantification of network parameters for CD07 (**C**) and CD19 (**D**) mutant and wild type lines. For CD07 bar graphs represent averaged recordings from days 41, 44, and 48. Each day contains five replicates for two isogenic pairs. For CD19, bar graphs represent averaged recordings from days 41 and 48. Each day contains five replicates for two isogenic pairs. Circles represent individual recordings. Student’s t-test was used to quantify statistical significance.

### *OTUD7A* LoF reduces expression of genes related to synaptic signaling and neuropsychiatric disorders

Having demonstrated the detrimental effects of *OTUD7A* LoF on synapse development and neuronal function in human iNs, we sought to explore the underlying molecular mechanism and its relevance to SZ and other neuropsychiatric disorders. We carried out RNA-sequencing (RNA-seq) and performed transcriptomic profiling of the isogenic CRISPR-edited iNs carrying the *OTUD7A* LoF mutation. Hierarchical clustering of genes with the most variable expression, i.e., top 500 differentially expressed genes (DEGs), revealed that RNA-seq samples could be clearly separated into two groups, the mutant iNs (from 2 different lines, each with 2 independent clones) and the wild type controls (2 different lines, each with 3 independent clones) (Figure 6A). We did not identify any DEGs at a false discovery rate (FDR) < 0.05; however, with a more permissive statistical cut-off (*p* < 0.05), we found 1,931 DEGs (992 downregulated, 939 upregulated) in mutant iNs compared to the wild type lines (Figure 6B and Table S2).

**Fig 6.**
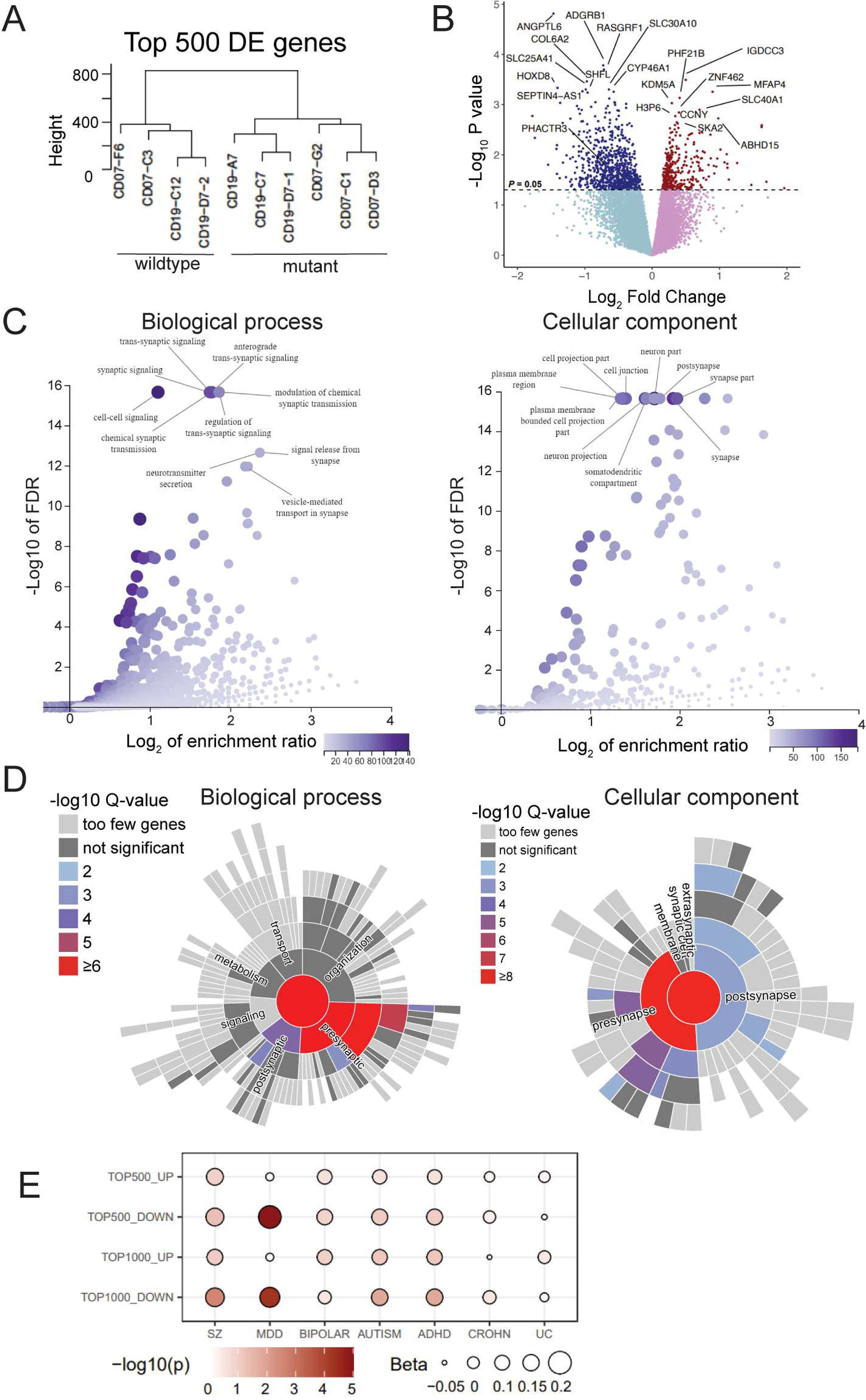
*OTUD7A* LoF reduces the expression of genes related to synaptic signaling and neuropsychiatric disorders. (A) Hierarchy cluster analysis of genes with the most variable expression, i.e., top 500 DEGs shows separations of RNA-seq samples into two groups, the mutant iNs (from 2 different lines, each with two independent clones) and the wild type (2 different lines, each with three independent clones). **(B)** Volcano plot of 1,931 DEGs in wild type and mutant lines (992 downregulated and 939 upregulated). Statistical cut off: *p* < 0.05. **(C)** GSEA using WEBGestalt shows that downregulated DEGs (*p* < 0.05) in the mutant iNs have strong enrichment (FDR ≤ 0.05) for genes associated with synapse, neuronal projection, neurotransmitter secretion, vesicle-mediated transport in synapse, and synaptic signaling. **(D)** SynGO analysis shows that downregulated DEGs (*p* < 0.05) in the mutant iNs have strong enrichment for biological process such as synaptic functions and synaptic vesicle release, and for cellular components of presynapse and postsynapse. **(E)** MAGMA analysis shows that the top-1,000 downregulated genes have significant enrichments for GWAS risk for major neuropsychiatric disorders. On the contrary, for the upregulated gene sets, no strong enrichment of GWAS risk for any neuropsychiatric disorders was found.

With the DEGs, we first examined whether the up- or downregulated genes were enriched for specific gene pathways relevant to our observed neuronal phenotypes associated with *OTUD7A* LoF. Gene set enrichment analysis (GSEA) using WEB-based GEne SeT AnaLysis Toolkit (WEBGestalt) (Liao et al., 2019) revealed that the downregulated DEGs (*p* < 0.05) in the mutant iNs showed strong enrichment (FDR < 0.05) for genes associated with synapse, neuronal projection, neurotransmitter secretion, vesicle-mediated transport in synapse, and synaptic signaling categories (Figure 6C and Table S3). However, the upregulated genes did not show any enrichment of genes related to neurodevelopment and synaptic function, but instead showed enrichment for genes related to cell cycle and DNA replication (Figure S5A and Table S3).

The observed specific enrichments for genes related to synapse and synaptic signaling among the downregulated, but not the upregulated genes, in the mutant iNs were further corroborated by a more focused Synaptic Gene Ontology (SynGO) analysis (Koopmans et al., 2019). Out of the 992 downregulated genes (*p* < 0.05) in the mutant iNs, 140 were mapped to unique SynGO annotated genes. Among these 140 genes, 113 were assigned to biological processes related to synaptic functions (enrichment *q*-value < 8.64 × 10^-11^) (Figure 6D), including 53 genes for presynaptic processes (enrichment *q*-value < 8.64 × 10^-11^) and 33 for postsynaptic processes (enrichment *q*-value < 6.58 × 10^-5^). 127 genes out of the 140 mapped unique SynGO annotated genes were assigned to presynaptic (enrichment *q*-value < 8.40 × 10^-10^) or postsynaptic (enrichment *q*-value< 2.04 × 10^-3^) cellular components (Figure 6D). However, for the upregulated genes, we did not observe enrichment for any biological process or cellular component related to synaptic function or synapse (Figure S5B). SynGO analysis using a more stringent statistical cut-off (*p* < 0.01) for selecting the down- and upregulated genes yielded similar results, with only downregulated DEGs showing significant enrichment for presynaptic (*q*-value < 5.31 × 10^-5^) or postsynaptic (*q*-value < 8.76 × 10^-3^) cellular components, and for biological processes such as presynaptic functions (*q*-value < 1.60 × 10^-9^) and synaptic vesicle cycle (*q*-value < 8.53 × 10^-8^). Altogether, the GSEAs suggest that *OTUD7A* LoF downregulated genes are involved in synapse and synaptic signaling, likely contributing to the altered neuronal phenotypes in the mutant iNs.

To further understand how *OTUD7A* LoF may act on the set of downregulated genes to impair synaptic development and neuronal function in iNs, we performed a Search Tool for the Retrieval of Interacting Genes/Proteins (STRING) (Szklarczyk et al., 2019) analysis of the protein-protein interaction (PPI) using all the downregulated DEGs (*p* < 0.01) together with OTUD7A. We reasoned that these downregulated genes were likely to form a PPI network including OTUD7A. Indeed, we found that these genes were strongly enriched for PPI (*p* < 1.0 × 10^-16^), in which OTUD7A was a peripheral network node. OTUD7A directly interacted with neuralized E3 ubiquitin protein ligase 1 (NEURL1) that further connected to a hub gene, sodium channel beta-3 subunit (SCN3B) (Figure S6A). It is noteworthy that NEURL1 is known to promote the growth of hippocampal dendritic spines and the increase of the GluA1 (Pavlopoulos et al., 2011), while SCN3B is known play a role in neuronal excitability (Cusdin et al., 2010; Oginsky et al., 2017). The PPI network was also enriched for gene ontology (GO)-terms related to synaptic function and nervous system development (Figure S6B), suggesting a possible mechanistic link between *OTUD7A* LoF and the observed neuronal phenotypic changes in the mutant iNs (Figures 4 and 5).

### *OTUD7A* LoF downregulated genes in iNs are relevant to SZ and other neuropsychiatric disorders

Finally, we examined whether *OTUD7A* LoF-induced transcriptional changes in iNs were relevant to SZ and other neuropsychiatric disorders. To test this, we performed Multi-marker Analysis of GenoMic Annotation (MAGMA) (de Leeuw et al., 2015) analysis to identify any enrichment of neuropsychiatric disorder GWAS signals in different sets of DEGs (top 500 or 1,000 hits, with later selection equivalent to the DEG *p*-value cut-off of 0.05) (Figure 6E). We found that the top 1,000 downregulated genes showed significant enrichment for GWAS risk loci of major neuropsychiatric disorders including SZ (Consortium et al., 2020) (*p* < 2.8 × 10^-3^), major depressive disorder (MDD) (Howard et al., 2019) (*p* < 4.8 × 10^-5^), autism spectrum disorder (ASD) (Grove et al., 2019) (*p* < 0.015), and attention deficit/hyperactivity disorder (ADHD) (Demontis et al., 2019) (*p* < 0.015), but not for bipolar disorder (Mullins et al., 2021) (Figure 6E). To a lesser extent, the top 500 downregulated genes also showed significant enrichment of GWAS risk loci for some of these neuropsychiatric disorders. However, for the upregulated gene sets, we did not find enrichment of GWAS risk loci for any neuropsychiatric disorders (Figure 6E). As expected, for the two “control” GWAS datasets (Crohn’s disease and ulcerative colitis) (Liu et al., 2015), we did not observe any significant enrichment in these DEG sets (Figure 6E). These results suggest that *OTUD7A* LoF down-regulated DEGs are specifically relevant to SZ and to some other neuropsychiatric disorders, which is congruent with our observed enrichment of GO-terms related to synapse and synaptic function only in downregulated DEGs (Figure 6C and D).

## DISCUSSION

The 15q13.3 deletion CNV has been found to be associated with various NDDs including SZ (Freedman et al., 2001; Gillentine and Schaaf, 2015; Helbig et al., 2009; Pagnamenta et al., 2009; Sharp et al., 2008; Shinawi et al., 2009; Stefansson et al., 2008; Ziats et al., 2016). A previously studied animal model suggested *Otud7a* in this CNV region as a plausible “driver” gene for the 15q13.3 deletion syndrome phenotypes (Yin et al., 2018). However, supporting evidence for the driver role of *OTUD7A* in the 15q13.3 deletion has been lacking. Here, our targeted resequencing of all SZ-associated CNVs identified *OTUD7A* LoF mutations present only in SZ cases, providing supportive human genetic evidence for the possible driver role of *OTUD7A.* Leveraging the rare *OTUD7A* LoF mutations identified in SZ cases, we further employed an isogenic CRISPR/Cas9 gene editing approach to engineer one of the *OTUD7A* LoF mutations (rs757148409/p.Arg89Ter) on different genetic backgrounds of healthy control donors and demonstrated that *OTUD7A* LoF impaired dendritic arborization and glutamatergic synapse development as well as population neural network activity in human iNs. We also found that the *OTUD7A* LoF-induced neurodevelopmental deficits were accompanied by transcriptome-wide downregulation of genes involved in synaptic function and genetically relevant to SZ and other neuropsychiatric disorders. Our findings thus support an important role of *OTUD7A* in the neurodevelopmental effects of the 15q13.3 deletion associated with SZ and other NDDs.

The cellular phenotypic effects of the SZ-associated 15q13.3 deletion have not been previously investigated in human neurons. Our observed reduction of dendritic complexities and glutamatergic synapse in hiPSC-derived excitatory iNs carrying the *OTUD7A* LoF mutation is consistent with what has been demonstrated in *Otud7a* knockout mice (Yin et al., 2018). More importantly, the *OTUD7A* LoF mutation identified in our MGS SZ case seemed to result in neural phenotypes relevant to SZ. SZ cases often show a reduction of cortical grey matter volume and thickness, as well as reduced functional cortical connectivity (Goghari et al., 2007; Harris, 1999; Lewis and Gonzalez-Burgos, 2008; Rapoport et al., 1999). The reductions in dendritic spine density on cortical pyramidal neurons observed in SZ postmortem brains are thought to directly contribute to brain imaging abnormalities in SZ cases (Goghari et al., 2007; Harms et al., 2010; Prasad et al., 2010; Rapoport et al., 1999). Reduced dendrite arborization has also been reported in some studies using hiPSC-derived neurons (Brennand et al., 2011; Flaherty et al., 2019; Wen et al., 2014). However, disease-relevant specific cellular phenotypes for neuropsychiatric disorders remain largely elusive. For instance, recent studies using hiPSC or mouse models found that SZ risk alleles were also associated with increased dendritic complexities and neuronal hyperexcitability (Blizinsky et al., 2016; Forrest et al., 2017; Schrode et al., 2019; Yi et al., 2016; Zhang et al., 2020). These studied SZ risk variants include both common variations (e.g., a GWAS SNP at the *MIR137* locus) (Forrest et al., 2017) with small effect sizes and rare risk alleles with high penetrance (e.g., 16p11 duplication or LoF of *SHANK3*) (Blizinsky et al., 2016; Yi et al., 2016). Such seemingly inconsistent cellular phenotypes may reflect pleiotropy of genetic risk effects across major psychiatric disorders (Ruderfer et al., 2018). In this regard, it is noteworthy that the *OTUD7A* LoF downregulated genes in iNs were found to be enriched for GWAS risk of not only SZ, but also some other major neuropsychiatric disorders (Figure 6E).

*OTUD7A* encodes a putative deubiquitinating enzyme. The *OTUD7A* LoF-induced neurodevelopmental deficiency observed in our study underscores an important role of ubiquitination or deubiquitination in 15q13.3 deletion-related neurodevelopmental or neuropsychiatric disorders. Interestingly, a recent study using hiPSC-derived neurons from patients with the 15q11-q13 duplication syndrome, a genetic form of autism, showed that *UBE3A,* a gene encoding an E3 ubiquitin ligase protein, may be the “driver” of the 15q11-q13 duplication syndrome (Fink et al., 2021). Other protein ubiquitination, deubiquitination, and degradation pathway genes have also been shown to carry excessive mutations in NDD cases. For instance, *E3 ubiquitin ligase cullin 3* (*CUL3*) is among the genes most confidently implicated 0 0.01) (Kaplanis et al., 2020; Matsuura et al., 1997; Sanders et al., 2015; Uddin et al., 2018). The balancing act of ubiquitinating and deubiquitinating enzymes has been shown to control neuronal fate determination, axonal pathfinding and synaptic communication, and plasticity (Todi and Paulson, 2011). Although we did not directly test the effect of the *OTUD7A* LoF mutation on ubiquitination/deubiquitination, the impaired ubiquitination and subsequent reduction of dendritic complexities and synaptic function represent a conceivable cellular mechanism underlying the genetic association of the 15q13.3 deletion with SZ and other NDDs.

How does *OTUD7A* LoF cause disorder-relevant neural phenotypes? This question may be addressed partially by our transcriptomic analysis of iNs carrying *OTUD7A* LoF, from which we found that downregulated genes were enriched for synapse development and function (Figure 6). Although the decreased expression of many synapse-related genes may simply reflect the secondary effect of the neuronal phenotype change rather than a “causal” factor, our results provided some mechanistic insights on how *OTUD7A* LoF works. First, our STRING (Szklarczyk et al., 2019) analysis indicated that the *OTUD7A* LoF downregulated genes tended to form into a PPI network linked with OTUD7A through a E3 ubiquitin-protein ligase (NEURL1), a gene that is known to promote the growth of hippocampal dendritic spines and the increase of the GluA1 (Pavlopoulos et al., 2011). Second, some of the *OTUD7A* LoF downregulated genes are strong candidates that may mediate the “causal” effect of *OTUD7A* LoF. For instance, the second most downregulated gene (log2_FC = -0.78, *p* = 7.3 × 10^-5^), *phosphatase and actin regulator 3* (*PHACTR3*) (Figure 6B, Table S2), encodes a neuron-enriched protein that regulates dendritic morphology (Miyata et al., 2020). Another plausible mediator and the third most downregulated gene (log2_FC = -0.64, *p* = 8.5 × 10^-5^), *Ras guanine nucleotide releasing factor 1* (*RASGRF1*) (Figure 6B, Table S2), serves as an NMDAR-dependent regulator of the ERK kinase pathway, facilitating neurotrophin-induced neurite outgrowth (Krapivinsky et al., 2003; Talebian et al., 2013). Third, the DNA-binding activity of OTUD7A conferred by its A20-like zinc finger protein domain may enable OTUD7A to act as a transcription factor (TF) or more plausibly, the *OTUD7A* LoF causes dysregulation of some specific TFs that further downregulate genes involved in synaptic function and neurodevelopment. In support of this hypothesis, our exploratory Enrichr (Kuleshov et al., 2016) analysis (Figure S7A,B) found that the *OTUD7A* LoF downregulated genes were enriched for multiple TF binding sites, of which the most enriched one, ZNF148 (also known as ZBP-89) (adjusted enrichment *p* = 5.8 × 10^-7^), could affect cellular growth and differentiation through its synergistic interactions with other TFs such as SP1 (Chupreta et al., 2007), which was also highly enriched in putative regulatory sequences of the *OTUD7A* LoF downregulated genes (adjusted enrichment *p* = 9.2 × 10^-6^).

It is intriguing that genes downregulated by *OTUD7A* LoF in iNs were found to be significantly enriched for GWAS risk of SZ and some other neuropsychiatric disorders (Figure 6E). This suggests that *OTUD7A* may partially drive clinical phenotypes of SZ and other neuropsychiatric disorders through acting on common GWAS risk genes. It has been shown that cases carrying rare and highly penetrant disease risk variants such as CNVs often have an excess burden of common GWAS risk alleles (Tansey et al., 2016). The field has just started to understand the interaction between individual polygenic risk scores (PRS) and highly penetrant rare risk variants such as the well-known 22q11 deletion (Cleynen et al., 2020). However, the mechanistic link between common genetic risk backgrounds (i.e., PRS) and rare disease risk variants has not been empirically tested. Interaction between rare and highly penetrant risk variants and PRS may also explain incomplete penetrance or phenotypic heterogeneity associated with rare disorder risk variants. Our observed enrichment of GWAS risk of SZ and some other neuropsychiatric disorders among genes downregulated by *OTUD7A* LoF thus provided empirical evidence supporting the possible interaction between common genetic risk backgrounds and rare disease risk variants, providing a foundation for studying how common genetic risk background may impact the phenotypic expressivity of 15q13.3 deletion.

Although our transcriptomic analyses suggested a possible mechanistic link between *OTUD7A* LoF and the impaired neuronal phenotypes, the profound transcriptomic effects of *OTUD7A* LoF in CRISPR/Cas9-engineered iNs may need to be interpreted cautiously. It has been a general challenge in the field that transcriptomic analysis of hiPSC-derived neurons, even for isogenic pairs of iPSC lines, can be confounded by line-to-line and/or clonal variations (Pak et al., 2021b). However, such limitations may be mitigated by our use of multiple lines/clones and the congruence between GO analysis of the DEGs and the neuronal phenotypic alterations between isogenic pairs of neurons. Other limitations of our study include: (1) We only studied *OTUD7A* LoF in hiPSC-derived excitatory neurons, while other cell types such as inhibitory neurons may also be important for *OTUD7A*. However, our choice of excitatory neurons was based on the observed higher expression of OTUD7A in excitatory neurons than in other cell types (Figure S2B,C). (2) Although the iNs used here represent a relatively homogenous cellular model, the two-dimensional (2D) cell culture has its intrinsic limitation in recapitulating *in vivo* neural phenotypes. Future studies using 3D brain organoids or assembloids (Paş recently demonstrated for modeling highly penetrant disease mutations (Kang et al., 2021; Khan et al., 2020), will better illuminate the neurodevelopmental and synaptic deficits associated with *OTUD7A* LoF mutations. (3) Among multiple genes in the 15q13.3 deletion, we only tested the functional effects of *OTUD7A* LoF; other genes may also contribute to the risk to SZ and other NDDs. Nevertheless, for the first time in human neurons, our study has illuminated a mechanistic link between *OTUD7A* LoF and neural phenotypes relevant to SZ and other NDDs, supporting a major role for *OTUD7A* in the 15q13.3 deletion syndrome.

## EXPERIMENTAL MODEL AND SUBJECT DETAILS

### hiPSC lines and cell culture

The hiPSC lines CD07 and CD19 (CD0000007 and CD0000019 respectively) used in CRISPR/Cas9 editing for genetic perturbation were derived at the Rutgers University Cell and DNA Repository (RUCDR)-NIMH Stem Cell Center. The sources cells for the hiPSC lines were cryopreserved lymphoblasts of the MGS collection (Sanders et al., 2008; Sanders et al., 2010; Shi et al., 2009). The hiPSC line was generated by using Sendai virus method to assure its being integration-free. The CD0000007 hiPSC line was from a 59 year old control male, and the CD0000019 hiPSC line was from a 41 year old control male. hiPSCs were cultured using a feeder-free method on matrigel (Thermofisher)-coated plate in mTeSR plus media (StemCell). Media were changed every other day and cells were passaged as clumps every 4-6 days using ReLeSR (StemCell). Confirmation of the absence of chromosomal abnormality was performed by eSNP-Karyotyping (Weissbein et al., 2016).

## METHOD DETAILS

### Targeted DNA sequencing, variant calling and quality control

A total of 1,963 SZ cases and 1,535 controls of European ancestry from the MGS collection (Sanders et al., 2008; Sanders et al., 2010; Shi et al., 2009) were selected for sequencing. DNA was extracted, prepared for PCR amplification, and amplified on the Raindance Thunderstorm system in batches of 96 samples. A total of 241 genes were targeted, covering 6 SZ-associated CNVs (Bassett et al., 2010; Levinson et al., 2011; Marshall et al., 2017; Szatkiewicz et al., 2014). The DNA samples were sequenced at Hudson Alpha using Illumina HiSeq 2500 in rapid run mode, with 2 x 250bp paired end reads, in batches of 384. We targeted high sequencing coverage (> 100×) to improve detection of rare variants. Sequence mapping and trimming were conducted using PEMapper (Johnston et al., 2017), against the hg38 human genome assembly.

Variants were called using PECaller (Johnston et al., 2017), using a repeat-masked target bed file to limit calls to the targeted and unique regions in hg38. The resulting .snp variant call file contained 3,418 samples (1,914 SZ cases, 1,500 controls, 4 with missing phenotypes) and 24,463 variants. This file was converted from the PECaller .snp format to the .vcf variant call format for annotation with Bystro (Kotlar et al., 2018). After sample exclusion because of apparent mislabeling or large deviation from the mean of statistics such as heterozygosity, sequencing completeness, transition/transversion ratio, Waterson’s theta, and ratio of heterozygotes/homozygotes, a total of 3,201 clean samples (1,779 SZ cases, 1,418 controls, 4 with missing phenotypes) remained. The final 3,201 samples were re-annotated in Bystro (Kotlar et al., 2018). Using Bystro’s search engine, variants were dropped if they had 1) missingness greater than 5%, or 2) allele frequencies different than those in gnomAD at a *p*- value < 2.4x10^-6^, unless those differing sites had 2 or fewer minor alleles (a maximum sample minor allele frequency of 4x10^-4^). This query was saved as a new annotation using the Bystro web application’s save feature, and further partitioned for statistical analysis.

### Variant filters and association test

Three variant sets were chosen for evaluation: 1) predicted deleterious variants, having a combined annotation dependent depletion (CADD) score > 15, 2) moderately rare deleterious variants, having a population allele frequency of 1x10^-2^ or less and a CADD > 15, and 3) rare deleterious variants, with a population allele frequency of < 1x10^-4^ and CADD > 15. The CADD threshold was chosen as it is the median score for missense and splicing variants. Population allele frequency estimates were based on the gnomAD non-Finnish European sample. SKAT-O optimal burden/variance-component test for per-gene and gene-region analyses (Lee et al., 2012). Sex, batch, and the top four principal components explaining genotype variance were included as covariates. For the gene-by-gene tests, *p*-values were corrected using the SKAT-O package’s parametric bootstrap method (Lee et al., 2012), adjusting for the number of genes included in those gene sets. Significance was evaluated at a 5% family-wise error rate.

### Sanger sequencing for mutation confirmation

To confirm mutations in SZ cases, DNA was isolated and Sanger sequencing was performed on three subjects with following IDs: 140-2257-001 (rs757148409/p.Arg89Ter), 145-1349-001 (c.1465_1473delAACGGCAAG/p.489-491(AspGlyLys)), and 148-2319-001 (c.406G>T/p.Glu136Ter). The region with a mutation was amplified by PCR using 10× PCR amplification buffer (Invitrogen), 10× PCR enhancer (Invitrogen), 50 mM MgSO_4_, 10 mM dNTPs, Forward and Reverse primers 8 µM each, 0.1 µl Hot Start Polymerase (Qiagen, 5 U/ µl), 20 ng/µl DNA template (95°C for 5 min, 39 cycles of [95°C for 30 sec, 55.4°C for 30 sec, 72°C for 1 min]). Then Shrimp Alkaline Phosphatase (SAP) reaction (total volume 10 µl) was performed with 10× SAP buffer (Affymetrix), 0.5 µl of SAP (Affymetrix), 0.1 µl *E. coli* exonuclease (Applied Biosystems, 10 U/ µl), 5 µl PCR product (37°C for 45 min, 95°C for 15 min). Sequence PCR (total volume 5 µl) was carried out with 1 µl Big dye (Applied Biosystems), 1 µl Forward primer, 5 µl SAP reaction (96°C for 1 min, 25 cycles of [96°C for 10 sec, 55°C for 5 sec, 60°C for 4 min]). Sequence PCR reaction was then purified: 40 µl 70% isopropanol was added and incubated for 20 min at room temperature, centrifuged at 3000 × g for 30 min, supernatant aspirated by inverting plate onto paper towel, pellet was rinsed by adding 150 µl 70% isopropanol, centrifuged at 2000 × g for 1 min, wash was discarded by inverting the plate onto paper towel, inverted plate with paper towel was centrifuged at 200 × g for 1 min. Sequencing was performed: samples were suspended in 15 µl formamide, centrifuged at 200 × g for 30 sec, denatured at 95°C for 2 min and immediately chilled on ice for 2 min, centrifuged at 2000 × g for 1 min. Samples were sequenced in 3730 DNA Analyzer with POP-7 Polymer (384) (Applied Biosystems) Primers for sequencing were synthesized at Integrated DNA Technologies (IDT) (Table S7).

### CRISPR/Cas9 editing of hiPSCs

CRISPR guide RNA (gRNA) sequences were designed using online tool (crispr.mit.edu), and we selected the gRNAs with highest score (specificity). The gRNAs were cloned into pSpCas9(BB)-2A-Puro vector (Addgene) for co-expression with Cas9 based on an established protocol (Ran et al., 2013). A day before transfection, we plated hiPSCs at 5 × 10^5^ cells on 60 mm plate using accutase in mTeSR plus media with 7.5 μM ROCK inhibitor. For transfection we combined 3 μg CRISPR/Cas9-gRNA construct with 3 μg ssODNs (1:1 ratio) in 200 μl Opti-MEM media (StemCell) in one tube. In the other tube we combined 12 μl Lipofectamine stem reagent (Invitrogen) (Lipofectamine:DNA ratio 2:1) in 200 μl Opti-MEM media. Then we combined two tubes and added the whole mixture to the hiPSCs. We started puromycin (0.4 μg/ml) selection at 24 h post transfection, reduced the puromycin to 0.2 μg/ml at 48 h post transfection, and withdrew it after 72 h of transfection. 14 days after transfection, resistant colonies were picked into 96-well plates and a small proportion of cells from each colony were used for DNA isolation using QuickExtract solution (Epicenter). The extracted DNAs were used for Sanger sequencing to verify editing. To obtain a pure heterozygous clone, each picked colony was subcloned at least two times and purity confirmed by Sanger sequencing. During subcloning cells were plated at a very low density on 60 mm plates (3 × 10^3^ cells). 14 days later, colonies were picked into 96-well plates and a small proportion of cells from each colony were used for DNA isolation using QuickExtract solution (Epicenter). The extracted DNAs were used for Sanger sequencing to verify the on-target and off-target editing. For checking off-target editing, primers were designed to amplify regions corresponding to the 5 top ranking predicted off-targets. Primers and oligos were synthesized at Integrated DNA Technologies (IDT) (Table S7).

### Quality control of the mutant hiPSC lines

To confirm the pluripotency of CRISPR-Cas9 edited hiPSC lines the cells were plated as clumps on coverslips. ∼7 days later when the hiPSC cells formed a colony, we fixed them in 4% PFA (Sigma) for 10 min at room temperature at Day 30 of neuronal differentiation. After 3 brief washes in PBS, cells were permeabilized with 1% Triton X-100 (Sigma) in PBS for 10 min at room temperature and blocked in 3% BSA (Thermofisher) and 0.3 % Triton X-100 in PBS at room temperature for 1 h. Then the samples were incubated with primary antibodies at room temperature for 1 h, followed by 3 PBS washes for 5 min each. The samples were then incubated with secondary antibodies at room temperature for 1 h. After another 3 PBS washes of 5 min each, samples were incubated in 0.5 μ, 6-diamidino-2-phenylindole) at room temperature for 10 min, washed 3 times with PBS 5 min each and mounted on glass slides using Prolong Diamond antifade reagent (Invitrogen). We prepared antibodies in PBS containing 3% BSA. The images were taken by Nikon ECLIPSE TE2000-U microscope. Primary antibodies used and their dilutions for incubation were: NANOG (Abcam Ab21624, 1:400), Oct-4 (Abcam Ab19857, 1:300), Tra-1-60 (Abcam Ab16288, 1:300). Secondary antibodies were: Alexa 488 donkey anti-mouse (A21202, 1:1000), Alexa 594 donkey anti-rabbit (A21207, 1:1000).

### NGN2 neuron differentiation and culturing

hiPSCs were dissociated into single cells by accutase and replated at 5 × 10^5^ cells per well on 6-well plates using mTeSR plus media with 5 μM ROCK inhibitor on Day (−1). On Day 0, cells were infected by 200 μl/well lentivirus cocktail containing 100 μl NGN2 virus and 100 μl rtTA virus (Zhang et al., 2013). The virus cocktail was prepared in mTeSR plus with 5 μM ROCK inhibitor. mTeSR plus media was switched to neuronal media consisting of Neurobasal media (Thermofisher), B27 supplement (Thermofisher), Glutamax (Thermofisher), and 2 μg/ml doxycycline (Sigma) on Day 1. After a two-day puromycin selection at 1 μg/ml on Day 2 and Day 3, it was withdrawn on Day 4. Cells were dissociated into single cells by accutase on Day 5 and replated on 6-well plates into 2 ml of neuronal culture media containing Neurobasal media, μg/ml doxycycline, 10 ng/ml BDNF (PeproTech), 10 ng/ml GDNF (PeproTech), and 10 ng/ml NT3 (PeproTech). Plates were coated with 0.1% PEI (Sigma) in 2× Borate buffer (Thermofisher) on Day 3 overnight, washed four times with sterile ddH_2_O on Day 4 and dried overnight. On day 6, 2 ml of neuronal media were added into each well. Neurons were fed every 3 days with neuronal culture media by removing 2 ml and adding 2 ml of fresh media. Doxycycline was withdrawn on Day 21.

For immunofluorescence staining, rat astrocytes (Sigma) were plated in a DMEM media (Thermofisher) with 10% FBS (Thermofisher) as a 100 μl were carefully removed and cells were plated as a 100 μl blob onto rat astrocytes at 0.5 × 10^4^ cells per well in neuronal culture media containing Neurobasal media, Glutamax, B27 supplement, 2 μg Neurobasal media, Glutamax, B27 supplement, 2 μg/ml doxycycline, 10 ng/ml BDNF, 10 ng/ml GDNF, 10 ng/ml NT3, and 5% FBS. On day 6, cells were fed with 400 μl of neuronal culture media. On day 6, 500 μl of were added into each well with half-volume of medium change every 3 days for continuous culturing. The FBS concentration in neuronal culture media was reduced to 1 % on day 14. Doxycycline was withdrawn on Day 21.

For morphology analyses, cells were plated similarly as for immunofluorescence staining with the following differences. On Day 28, 1 μg GFP plasmid and 2 μl Lipofectamine 2000 (Thermofisher) with DNA:reagent ratio 1:2 were added into 25 μl of DMEM media in separate with DNA:reagent ratio 1:2 were added into 25 μl of DMEM media in separatetubes and incubated for 5 min at room temperature. Then, GFP plasmid was combined with Lipofectamine reagent and incubated for another 20 min at room temperature. After that, the transformation mixture was added to the cells. Neurons were then fixed in 4% paraformaldehyde (PFA) containing 4% sucrose at room temperature for 15 min on Day 30.

### Immunofluorescence of NGN2 neurons

For routine characterization of iNs, neurons were fixed and stained as described in Quality control of the mutant hiPSC lines. The images were taken by Nikon ECLIPSE TE2000-U microscope. Primary antibodies used and their dilutions for incubation were vGlut1 (Synaptic Systems 135311, 1:300), MAP2 (Synaptic Systems 188011, 1:700), and HuNu (Millipore MAB 1281, 1:300). Secondary antibodies were Alexa 488 donkey anti-rabbit (A21206, 1:1000) and Alexa 594 donkey anti-mouse (A21203, 1:1000).

For immunostaining of iNs in dendritic and synaptic analyses, neurons were fixed at Day 30 (2 days after transfection with GFP) using 3.7 % formaldehyde (Sigma) in 4% sucrose/PBS for 15 min at room temperature. After 3 brief washes in PBS, neurons were permeabilized and blocked simultaneously in PBS containing 0.1% Triton and 5% normal goat serum (Jackson Immunoresearch) for 1 h at room temperature followed by incubation with primary antibodies overnight at 4°C. Neurons were washed 3 times in PBS 5 min each and incubated with secondary antibodies at room temperature for 1 h followed by another 3 washes in PBS of 5 min each. Coverslips with immunostained neurons were incubated in 0.5 μg/ml DAPI at room temperature for 10 min, washed 3 times with PBS for 5 min each, and mounted on glass slides using Prolong Gold antifade reagent (Invitrogen). Primary and secondary antibodies were diluted in PBS containing 5% normal goat serum. Neurons from the same differentiation experiment were fixed and stained at the same time with identical antibody dilutions. Primary antibodies used and their dilutions for incubation were GFP (Abcam ab13970, 1:10,000), GluA1 (Millipore AB1504, 1:300), and PSD-95 (NeuroMab clone K28/43, 1:1000). Secondary antibodies were Alexa 488 goat anti-chicken (A32931, 1:1000), Alexa 568 donkey anti-mouse (A11031, 1:1000), and Alexa 647 goat anti-rabbit (A32733, 1:1000).

### Dendritic complexity analysis

To quantify dendritic complexity, GFP-transfected neurons were imaged on a Nikon C2+ confocal microscope with NIS-Elements software. At least thirty neurons were imaged for each isogenic pair. Neurons were only imaged if a visible axon and at least one defined primary dendrite could be identified. Following acquisition, dendrites were traced and images binarized in ImageJ (National Institutes of Health, Bethesda, MD). For each image, the axon was identified and eliminated from quantification. Sholl analysis was performed using the ‘Sholl analysis’ plugin for ImageJ. The center of the soma was defined manually, and concentric circles spaced 10 μm apart were used to quantify number of intersections over distance in μm (dendritic complexity) in all neurons. Dendrite length was quantified by drawing a line along the dendrite and determining the length of this line in μm. All analyses were performed blind to genotypes.

### Puncta analysis

Images of GFP-transfected and immunostained neurons were acquired using a 63× oil immersion objective lens on a Nikon C2+ confocal microscope with NIS-Elements software. All images were acquired using identical settings for each channel to allow comparison between groups. For the images presented in Figure 4, each channel was adjusted for brightness and contrast and then merged to create the final composite images. Images were otherwise unaltered. Dendrites were imaged on multiple z-planes to capture full dendritic branches, and maximum intensity projections of z stacks created in ImageJ were used for downstream analyses. Five to ten neurons from multiple coverslips were imaged for each isogenic pair. Puncta were analyzed using the ‘Analyze Particle’ function in ImageJ. Regions of interest (ROI) on the dendrite (∼80 μm) were selected using the GFP-filled cell as an outline. A set threshold was defined for each experiment and applied identically to all conditions so that only puncta above diffuse background staining were considered in the analysis. The puncta area and intensity were recorded. PSD-95/GluA1 or colocalized puncta density was calculated as number of PSD-95/GluA1 puncta or colocalized puncta divided by area (puncta/µm^2^).

### MEA analysis

iNs at day 28 were replated on 24-well MEA plates (vendor) at 8 × 10^4^ cells per well along with rat astrocytes at 1.2× 10^4^ cells per well in neuronal culture media containing Neurobasal media, Glutamax, B27 supplement, 2 μg/ml doxycycline, 10 ng/ml BDNF, 10 ng/ml GDNF, 10 ng/ml NT3, and 1% FBS. Cell suspension was added as 10 μl blob which just covered the electrode area. Cells were fed with 500 μl of neuronal culture media 1 h later. Two days before replating plates were coated with 0.1% PEI in 2× Borate buffer as 10 μl blob. Sterile ddH2O was added in outer rim of the plate to prevent evaporation. Next day the plate was washed four times with sterile ddH_2_O and dried overnight. On day 29, cells were fed with 400 μl of neuronal media. On day 29, 500 μl of neuronal culture media were added into each well with half-volume of medium change every 3 days for continuous culturing. We started recording neuronal activity using Maestro Edge multiwell microelectrode array (Axion Biosystems) on day 34 every three days until day 60. Time of recording was 10 min. Media was always changed a day before recording to make the recordings consistent.

### Electrophysiology

Cells were transferred, on a cover slip, to a recording chamber and superfused with artificial cerebrospinal fluid (aCSF: 125 NaCl mM, 2.5 KCl mM, 1 MgCl_2_ mM, 2.5 CaCl_2_ mM, 20 glucose, 1 NaH_2_PO_4_ mM, 25 NaHCO_3_ mM and 1 mM ascorbic acid, bubbled continuously with 95% O2 / 5% CO_2_; perfused at 2 ml/min). Cells were visualized under infrared illumination using an upright Olympus BX51 microscope. Whole-cell voltage-clamp and current-clamp recordings used an Axopatch 200B amplifier, a Digidata 1200 interface, and pClamp 8 (Molecular Devices), and data were filtered at 1 kHz and digitized at 5 kHz, Vm = -70 mV. Borosilicate electrodes were pulled to a resistance of 3-7 MΩ and contained a recording solution (in mM): 154 K-gluconate, 1 KCl, 1 EGTA, 10 HEPES, 10 glucose, and 5 ATP, pH 7.4 with KOH. Series resistance was <20 MΩ. For measurement of cell responses to hyperpolarization and depolarization, recordings were conducted in current-clamp mode, and consisted of injection of a series of currents, via the recording pipette, ranging from -100pA to +200 pA. To obtain evoked synaptic responses, a concentric bipolar stimulating electrode was placed in the vicinity of the recorded cells, with recordings carried out in voltage-clamp mode. Excitatory synaptic responses to pairs of stimulation currents, using stimulus intensities ranging from 100 to 1,000 μA, and an interstimulus interval of 50 ms, were obtained. While we did not identify the nature of the excitatory responses, paired-pulse stimulation, at more depolarized potentials, did not reveal currents that might be identified as GABAergic, indicating that the evoked synaptic responses observed by us at -70 mV were likely excitatory and glutamatergic in nature.

### RNA isolation and RNA-seq

Cells from hiPSCs and iNs cultures were dissociated with accutase and washed with PBS. Then, cells were lysed in RLT plus buffer, and total RNA extracted with the RNeasy Plus Kit (Qiagen). cDNA was reverse transcribed from RNA with a high-capacity cDNA reverse transcription kit (Applied Biosystems).

The RNA sample was then cleaned with RNeasy MinElute Cleanup Kit (Qiagen) and subjected to RNA-seq at GENEWIZ (San Diego, CA). RNA-seq files were provided in the format of 2×150 bp paired-end (∼25 M reads per sample) fastq files from GENEWIZ. Briefly, cDNA libraries were prepared using the polyA selection method with Illumina HiSeq kit and customized adapters. No Trimmomatic-based adapter-trimming was performed as the sequencing facility had performed pre-cleaning on raw reads. 20∼30 M paired-reads were recovered from each sample at the rate of ∼350 M raw read-pairs per lane on an Illumina HiSeq 2000. Raw files were subsequently mapped to human hg38 genome (GRCh38p7.13) using STAR v2.7.0 and GENCODE v35 annotation file.

### Gene expression analysis by qPCR

For qPCR, reverse-transcription was performed using ThermoFisher High-capacity RNA-to-cDNA reverse transcription kit (Applied Biosystems, 4366596) with random hexamers according to the manufacturer’s protocol. Briefly, 300 ng of total RNA was used for each 20 µl RT reaction. The reaction products were then diluted with 80 µl of RNase-free water for qPCR analysis. qPCR was performed using TaqMan Universal PCR Master Mix (Applied Biosystems, 4364338) on a Roche 480 II instrument, using gene-specific FAM-labelled TaqMan probe (Hs00370128_m1) for detecting gene expression. Three technical replicates for each isogenic pair were included in qPCR. Relative gene expression was calculated as described (Forrest et al., 2017) with GAPDH as control.

### Western blotting

iNs at day 30 were dissociated with accutase, washed in PBS, spun down 5 min at 250×g and resuspended in 150 μl lysis buffer, which consisted of Phosphosafe buffer (EMD Millipore) with protease inhibitor (Roche) and PhosphoSTOP tablet (Roche). Cells were homogenized by pipetting up and down 20 times with P1000μl tips. The lysate then was transferred to QiaShredder (Qiagen) and spun down for 15 min at 14000 rpm at 4°C. The Pierce BCA protein assay (Thermofisher) was performed to determine protein concentration in the lysate. Protein sample was prepared by incubating 50 μg of protein lysate with 4× sample buffer (10% BME in sample was prepared by incubating 50 μg of protein lysate with 4× sample buffer (10% BME in4× Laemmlie Buffer). Protein sample was incubated at 98°C for 5 min and ran on Bio-Rad TGX Stain-Free Precast Gel (4-15%) in Tris/Glycine/SDS running buffer (BioRad). Then the gel was transferred onto a 0.45 μm PVDF membrane (Thermofisher). Before transfer the membrane was activated with methanol for 1 min and washed in anode buffer (15% methanol in Tris/CAPS buffer). Protein membrane was cut at ∼75kDa so it could be incubated with two different primary antibodies at the same time. After that the protein membrane was blocked with 5% skim milk (Thermofisher) in TBS-T wash buffer (Sigma) for 1 h at room temperature. OTUD7A and β primary antibodies were added into blocking buffer and the protein membrane was incubated at 4°C overnight followed by washing 3 times for 15 min with wash buffer. Secondary antibodies were added into blocking buffer and the protein membrane was incubated for 1 h at room temperature followed by washing 3 times for 15 min with wash buffer. Substrate (Millipore) was added to the protein membrane and the image was developed. To incubate with PSD-95 primary antibodies, the protein membrane was stripped in stripping buffer (Thermofisher) for 15 min in room temperature followed by 15 min wash and 30 min blocking. PSD-95 primary and secondary antibodies were applied as was described above followed by developing the membrane. Primary antibodies used and their dilutions for incubation were rabbit OTUD7A (Sigma, 1:1000), mouse PSD-95 (NeuroMab clone K28/43, 1:5,000), and mouse b-actin (Millipore, 1:20,000). Secondary antibodies were anti-rabbit-HRP (Cell Signaling Technology, 1:20,000) and anti-mouse-HRP (Cell Signaling Technology, 1:50,000 for PSD-95 and 1:100,000 for b-actin).

### RNA-seq data analysis

RNA-seq files were provided in the format of 2×150 bp paired-end (∼25 M reads per sample) fastq files from GENEWIZ. Briefly, cDNA libraries were prepared using the polyA selection method with Illumina HiSeq kit and customized adapters. No Trimmomatic-based adapter-trimming was performed as the sequencing facility had performed pre-cleaning on raw reads. 20∼30 M paired-reads were recovered from each sample at the rate of ∼350 M raw read-pairs per lane on an Illumina HiSeq 2000. Raw files were subsequently mapped to human hg38 genome (GRCh38p7.13) using STAR v2.7.0 and GENCODE v35 annotation file with the following parameters: STAR --bamRemoveDuplicatesType UniqueIdentical --outSAMtype BAM SortedByCoordinate --quantMode GeneCounts --outSAMattrIHstart 0 --outSAMstrandField intronMotif --outSAMmultNmax 1 --outFilterIntronMotifs RemoveNoncanonical -- outBAMsortingBinsN 40 --outFilterMultimapNmax 1 --outFilterMismatchNmax 2 -- outSJfilterReads Unique --alignSoftClipAtReferenceEnds No --outFilterScoreMinOverLread 0.30 --outFilterMatchNminOverLread 0.30. DGE analysis was performed using EdgeR by fitting the dataset into a gene-wise negative binomial generalized linear model with quasi-likelihood as glmQLFit() and evaluated by glmQLFTest(). After using the cut-off parameters mentioned above in STAR and further removing PCR and optical duplicates using Picard MarkDuplicates, the output BAM file has an overall unique alignment rate of > 65%.

### MAGMA analysis

We performed MAGMA analysis using MAGMA version 1.09 (Sey et al., 2020) to evaluate the enrichment for the GWAS risk of several mental disorders, as well as two related GWAS sets (Crohn’s disease and Ulcerative colitis) as the control set. Specifically, we first started by compiling the MAGMA-required gene annotation data files using a GRCh37/hg19 vcf file. With the gene-SNP annotation file, we then performed gene-level analysis on SNP P-values using the reference SNP data of 1,000 Genomes European panel (g1000_eur, --bfile) and the pre- computed SNP P-values from the GWAS data sets mentioned. The sample size (ncol=) were also either directly taken from the column of sample sizes per SNP column of the datasets or extracted from the affiliated README data. Subsequently, the result files (--gene-results) from the gene-level analysis were read in for competitive gene-set analysis (--set-annot), where we used default setting (‘correct=all’) to control for gene sizes in the number of SNPs and the gene density (a measure of within-gene LD). The gene-set analysis produced the output files (gsa.out) with competitive gene-set analysis results that contained the effect size (BETA) and the statistical significance of the enrichment of each gene set (top 500/1000 up/down genes) for each GWAS disease set.

### Gene set enrichment analysis

GSEA was performed using WEB-based (GEne SeT AnaLysis Toolkit) (Liao et al., 2019) to test downregulated DEGs in the mutant iNs for enrichments of GO terms. STRING 11.0 PPI network analysis (Szklarczyk et al., 2019) was conducted on the using all the downregulated DEGs together with OTUD7A.

### SynGo analysis

Synaptic GO analysis was used using SynGo (Koopmans et al., 2019) to test for specific enrichments for genes related to synapse and synaptic signaling among the downregulated DEGs in the mutant iNs.

### Statistical analyses

For Sholl analysis and MEA analysis, a two-way repeated-measures ANOVA was performed to quantify overall significant difference between mutant and wild type lines. For all other analyses, an unpaired Student’s t-test was used. All analyses were performed blinded to genotypes.

## SUPPLEMENTAL INFORMATION

Supplemental information can be found online.

## ACKNOWLEDGMENTS

MGS includes P. V. Gejman, A. R. Sanders, J. Duan (NorthShore University HealthSystem, and University of Chicago, IL, USA), D. F. Levinson (Stanford University, CA, USA), J. Shi (National Cancer Institute, MD, USA), N. G. Buccola (Louisiana State University Health Sciences Center, LA, USA), B. J. Mowry (Queensland Centre for Mental Health Research, Brisbane and Queensland Brain Institute, The University of Queensland, Australia), R. Freedman, A. Olincy (University of Colorado Denver, CO, USA), F. Amin (Atlanta Veterans Affairs Medical Center and Emory University, GA, USA), D. W. Black (University of Iowa Carver College of Medicine, IA, USA), J. M. Silverman (Mount Sinai School of Medicine, NY, USA), W. F. Byerley (University of California at San Francisco, CA, USA), C. R. Cloninger, D. M. Svrakic (Washington University, MO, USA). We thank the study participants of MGS. MGS was mainly supported by R01MH059571, R01MH081800, and U01MH079469 (to P.V.G.); and other NIH grants for other MGS sites (R01MH067257 to N.G.B., R01MH059588 to B.J.M., R01MH059565 to R.F., R01MH059587 to F.A., R01MH060870 to W.F.B., R01MH059566 to D.W.B., R01MH059586 to J.M.S., R01MH061675 to D.F.L., R01MH060879 to C.R.C., U01MH046276 to C.R.C., and U01MH079470 to D.F.L). We thank the PGC-SZ group for providing early access of wave-3 SZ (PGC-SZ3) GWAS summary statistics. We also thank Rutgers University Cell and DNA repository (RUCDR; 2U24MH068457) for producing hiPSC lines from the MGS cohort. Funding was provided by NIH grants: R01MH100915 (to P.V.G. and A.R.S.); R01MH100917 (to J.G.M and S.T.W); and R01MH106575 and R01MH116281 (to J.D.).

## AUTHOR CONTRIBUTIONS

A.K. (Kozlova) designed and performed the main experiments and data analyses and wrote the manuscript. S.Z. performed transcriptomic data analyses and wrote the manuscript. B.J. and H.Z. helped with hiPSC editing and neuronal differentiation. S.S. performed the MAGMA analysis. J.M. performed the patch-clamp experiment. M.P.F helped with neural dendritic and synaptic analyses. Z.P.P. helped with neuronal differentiation and neural phenotypic data interpretation. A.K. (Koltar) performed analysis of the targeted sequencing data. D.J.C., M.P.E., and M.E.Z provided bioinformatic and statistical guidance and helped with interpretation of sequencing results. A.R.S., S.T.W (deceased), P.V.G., and J.G.M. conceived the study, provided the sequencing data, and wrote/edited the manuscript. J.D. conceived the study, designed and supervised the experiments and data analyses, and wrote the manuscript.

## Web resources

The web links for the tools used in this study are: UCSC human genome browser (genome.ucsc.edu/cgi-bin/hgGateway); ClinVar (www.ncbi.nlm.nih.gov/clinvar/); SCHEMA- Schizophrenia exome meta-analysis consortium(schema.broadinstitute.org); gnomAD browser (gnomad.broadinstitute.org); WEB-based GEne SeT AnaLysis Toolkit (WEBGestalt) (www.webgestalt.org); Synaptic Gene Ontology (SynGO) (www.syngoportal.org); Cell type- specific single cell RNA-seq data in Allen Brain Map (portal.brain-map.org/atlases-and-data/rnaseq); String PPI network analysis (string-db.org); MAGMA gene set analysis (ctg.cncr.nl/software/magma).

## Data and code availability

RNA-seq data are available at NCBI GEO under accession number GSE189614. Codes for RNA-seq data analysis and for MAGMA GSEA are available upon request.

---

**Fig. S1:**
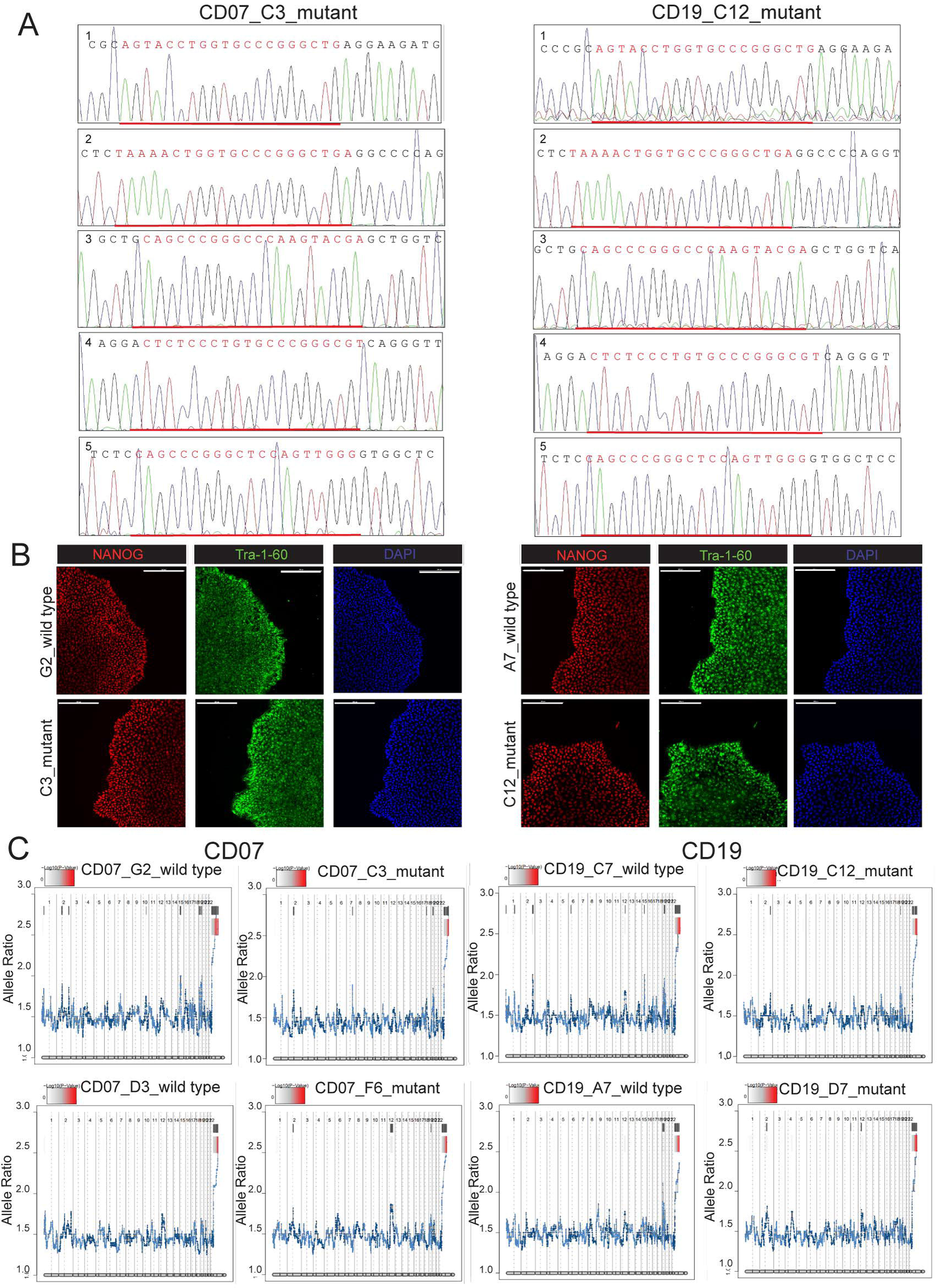
C**h**aracterization **of CRISPR/Cas9 engineered hiPSC lines and differentiated NGN2-induced excitatory neurons (iNs). (A)** The isogenic pairs of CRISPR-edited and wild type hiPSCs were sequenced to test for editing at five off-target genomic sites (underlined with red color). Representative sequences for CD07 and CD19 wild type are shown. **(B)** The isogenic pairs of CRISPR-edited and wild type hiPSCs were characterized for pluripotency by immunofluorescence staining (TRA-1-60+/NANOG+). Scale bar: 200 µm. **(C)** eSNP-karyotyping (using RNA-seq data) of isogenic CD07 (left panel) and CD19 (right panel) iNs to confirm the absence of chromosome abnormality. Shown are moving average plots of RNA-seq intensities of called SNPs along the chromosomes. No chromosome aneuploidy in hiPSC lines or iN-Glut neurons were identified. Color bars represent the -log_10_ value FDR-calibrated *p*-value.

**Fig S2:**
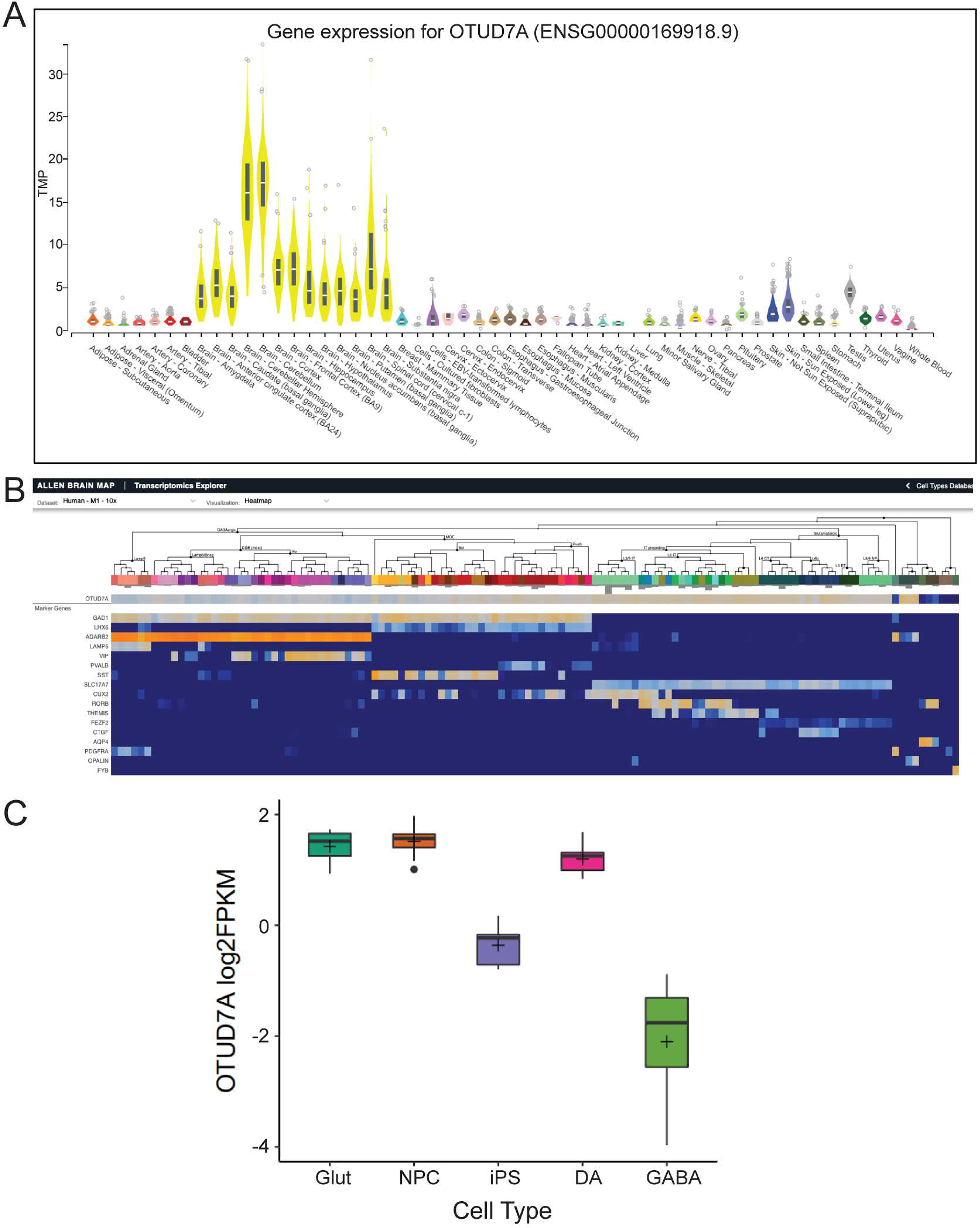
Brain-specific and hiPSC-derived neuronal cell type-specific expression of *OTUD7A*. (A) Gene expression data from the GTEx database showing that *OTUD7A* is predominantly expressed in the brain. **(B)** Hierarchical clustering of gene expression data in different neuronal cell types from the Allen Brain Map showing *OTUD7A* expression has a strong expression profile in glutamatergic neurons. **(C)** RNA-seq data from four different neuronal cell types plus hiPSC showing that *OTUD7A* has a strong expression in glutamatergic neurons, dopaminergic neurons, and NPCs. Quantification was measured as log_2_RPKM (Zhang et al, 2020).

**Fig S3:**
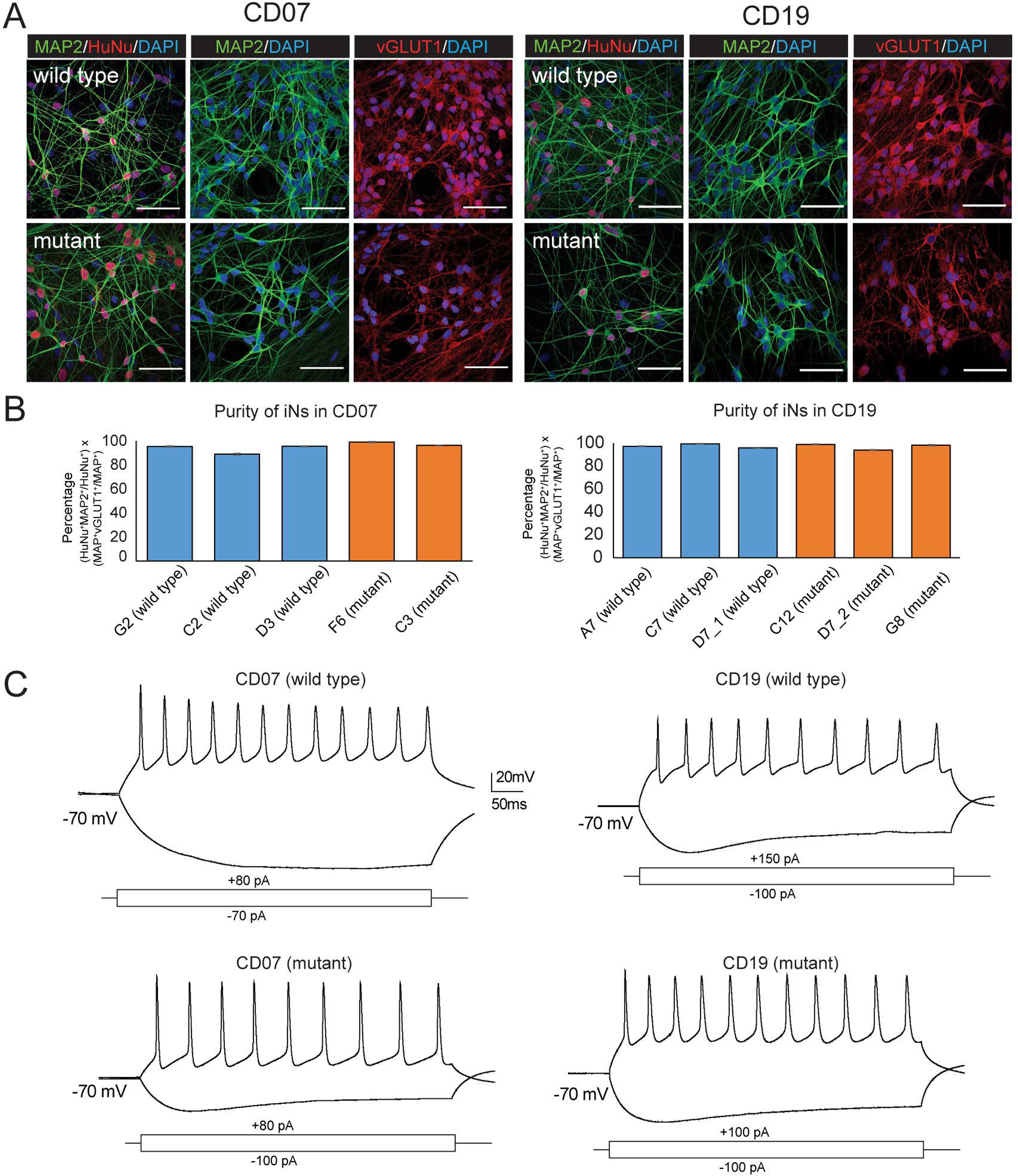
Purity of NGN2-induced neurons. (A) NGN2-induced neurons were stained for MAP2 and VGLUT1 to estimate their purity. Immunofluorescence images showing CD07 (left panel) and CD19 (right panel) isogenic iNs (MAP2+/VGLUT1+). Scale bar: 50 µm. **(B)** Percentage of neuronal cells that are stained positive for MAP2 and VGLUT1 for CD07 and CD19 lines. **(C)** Under whole-cell current clamp recording conditions, injection of negative current in CD07 (left) and CD19 (right) iNs, via the recording pipette, revealed hyperpolarization and voltage ‘sag’, while action potential trains were observed in response to positive current injection. In addition to triggering action potentials, the action potential firing rate response was proportional to the amount of positive current injected, and in some cases action potentials were preceded by pacemaker potentials and followed by after-hyperpolarization. All cells were held at -70mV, prior to current injection.

**Fig S4.**
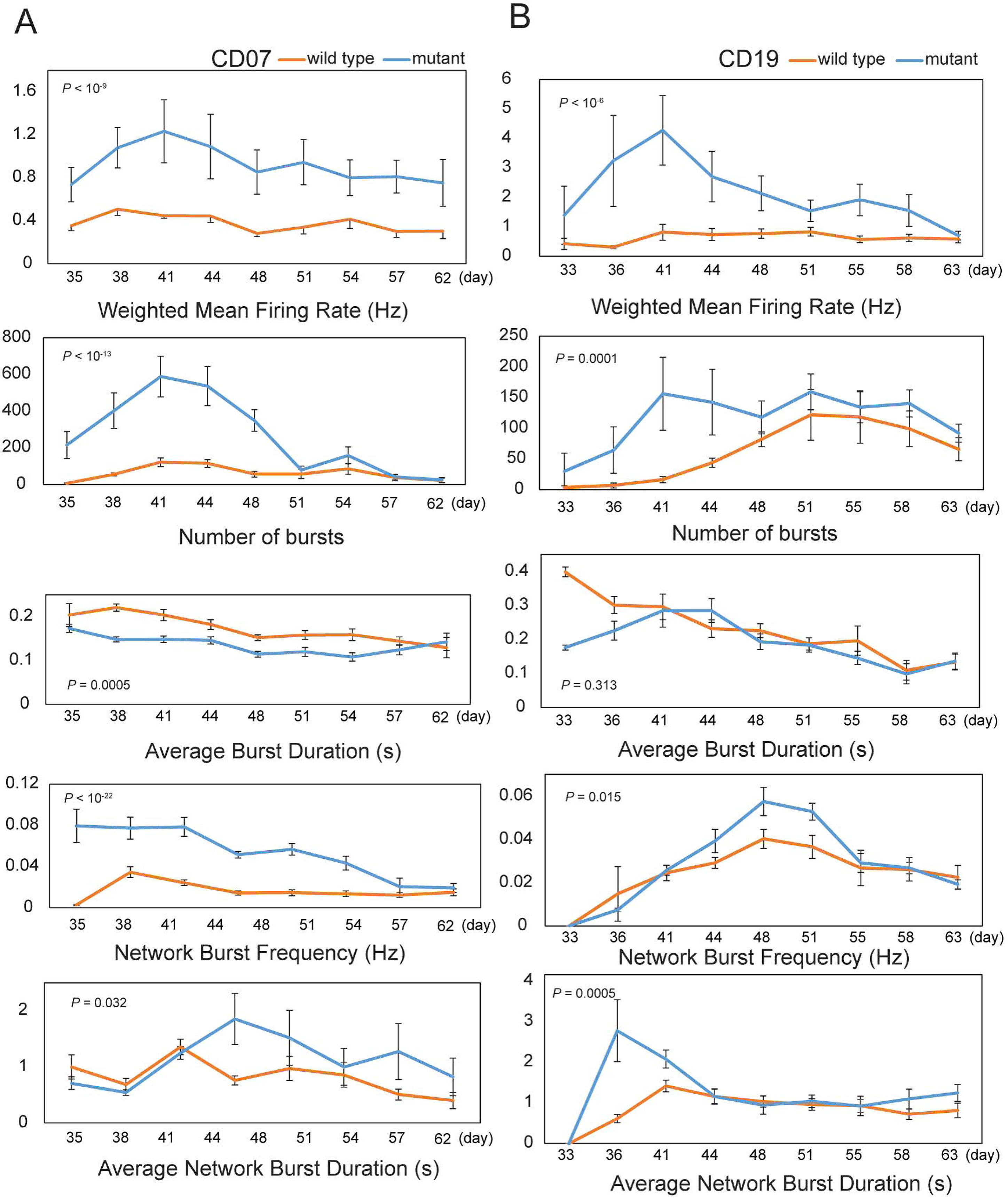
MEA analysis of *OTUD7A* mutant neurons shows reduced network activity. Quantification of network parameters as indicated in mutant CD07 (**A**) and CD19 (**B**) iNs compared to wild type from days 35 to 62 (CD07) and from days 33 to 63 (CD19) after induction. At each day, recordings were averaged among two isogenic pairs (five replicates for each clone) for both lines. Error bars are SEM. Two-way repeated-measures ANOVA was used to quantify statistical significance between edited and wild type lines.

**Fig S5.**
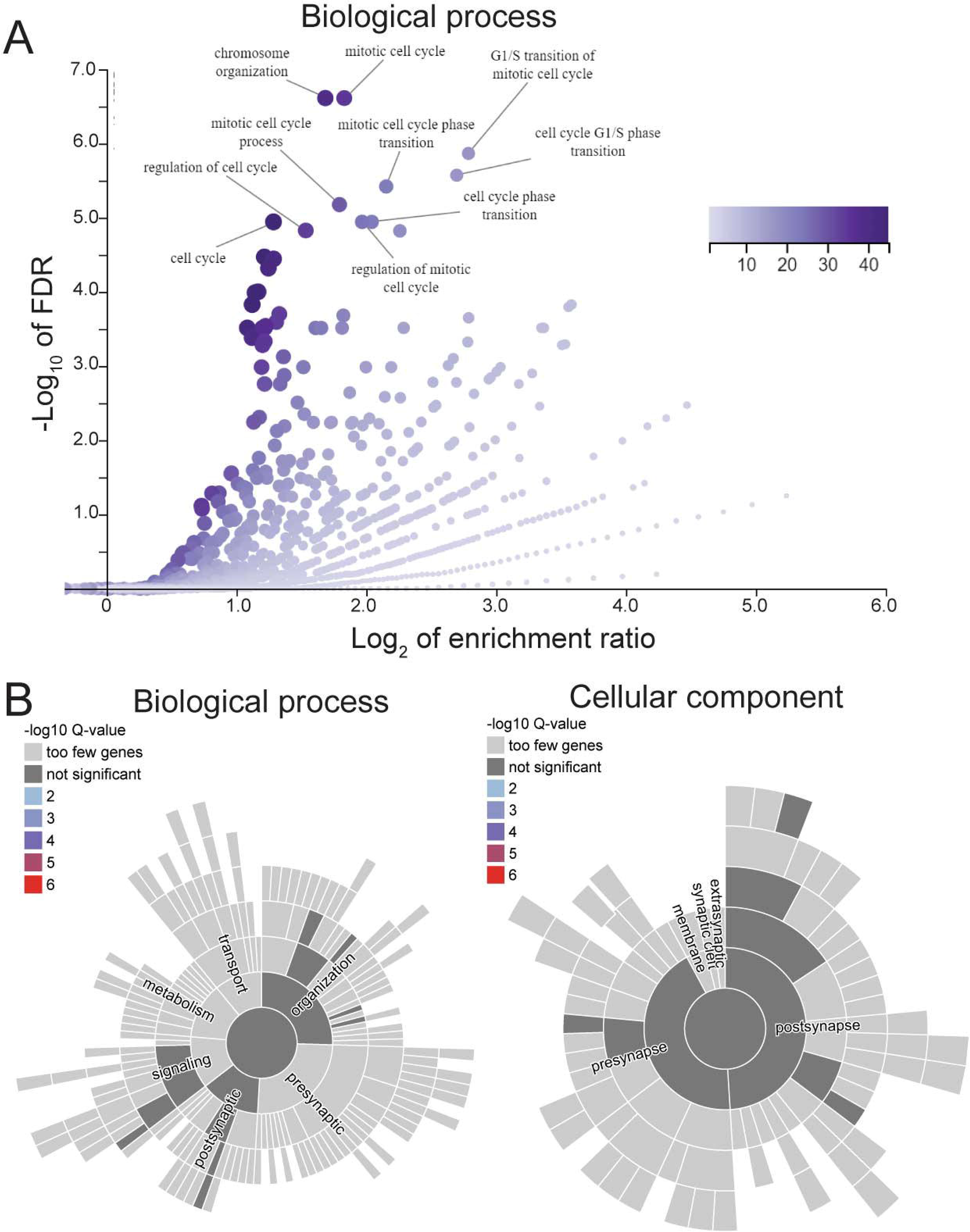
*OTUD7A* LoF elevates the expression of genes related to the cell cycle and DNA replication. (A) GSEA using WEBGestalt shows that upregulated DEGs (*p* < 0.05) showed enrichment for genes related to cell cycle and DNA replication. **(B)** SynGO analysis shows that upregulated DEGs (*p* < 0.05) did not show enrichment for any biological process or cellular component related to synaptic function or synapse.

**Fig S6.**
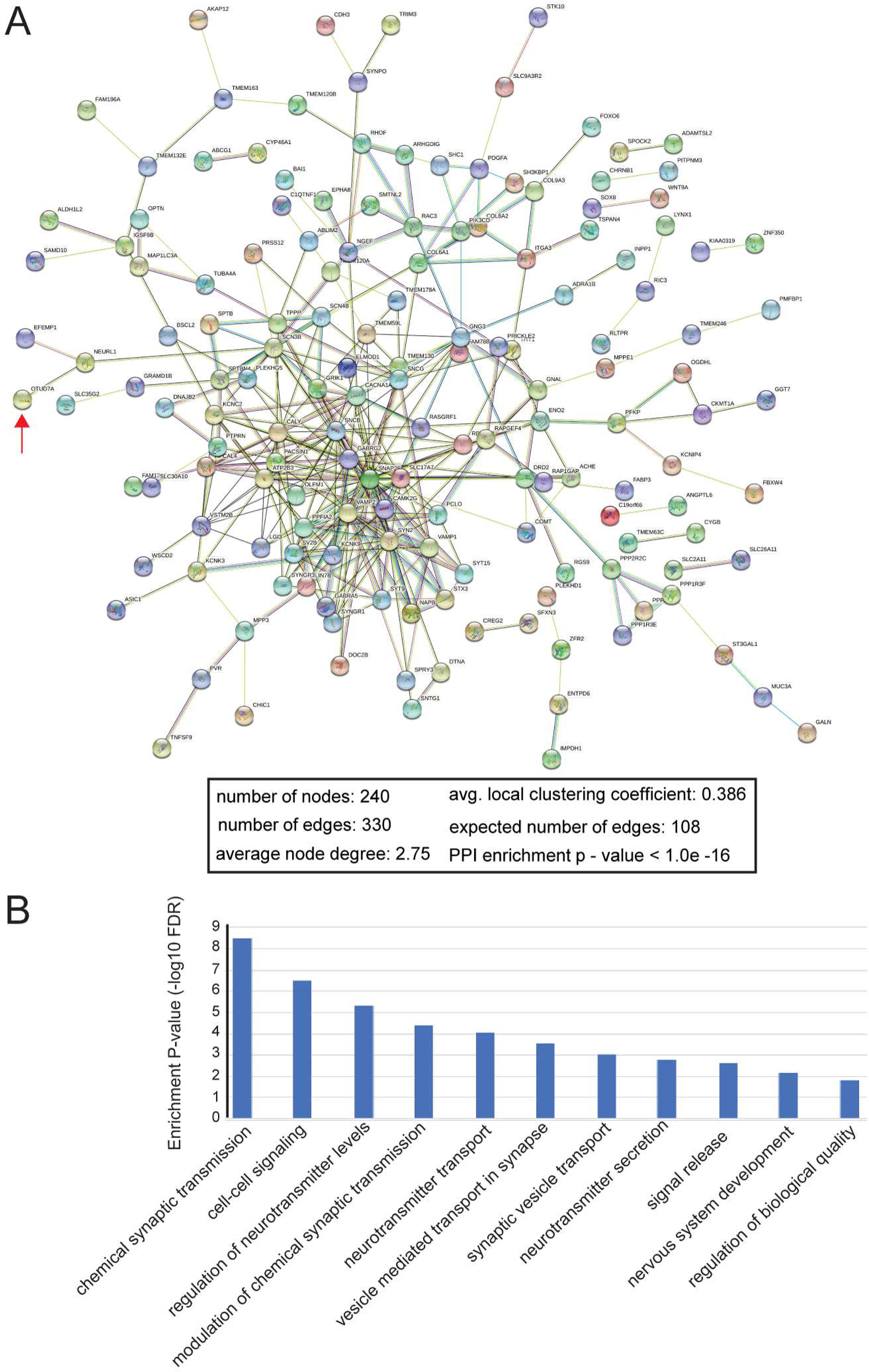
*OTUD7A* LoF may result in neuronal phenotypic changes in the mutant iNs. (A) STRING analysis shows that the downregulated DEGs (*p* < 0.01) together with OTUD7A form a PPI network. OTUD7A directly interacts with NEURL1 that further connects to SCN3B. OTUD7A is indicated with a red arrow. **(B)** Enrichment for GO-terms related to synaptic function and nervous system development in the OTUD7A PPI network.

**Fig S7.**
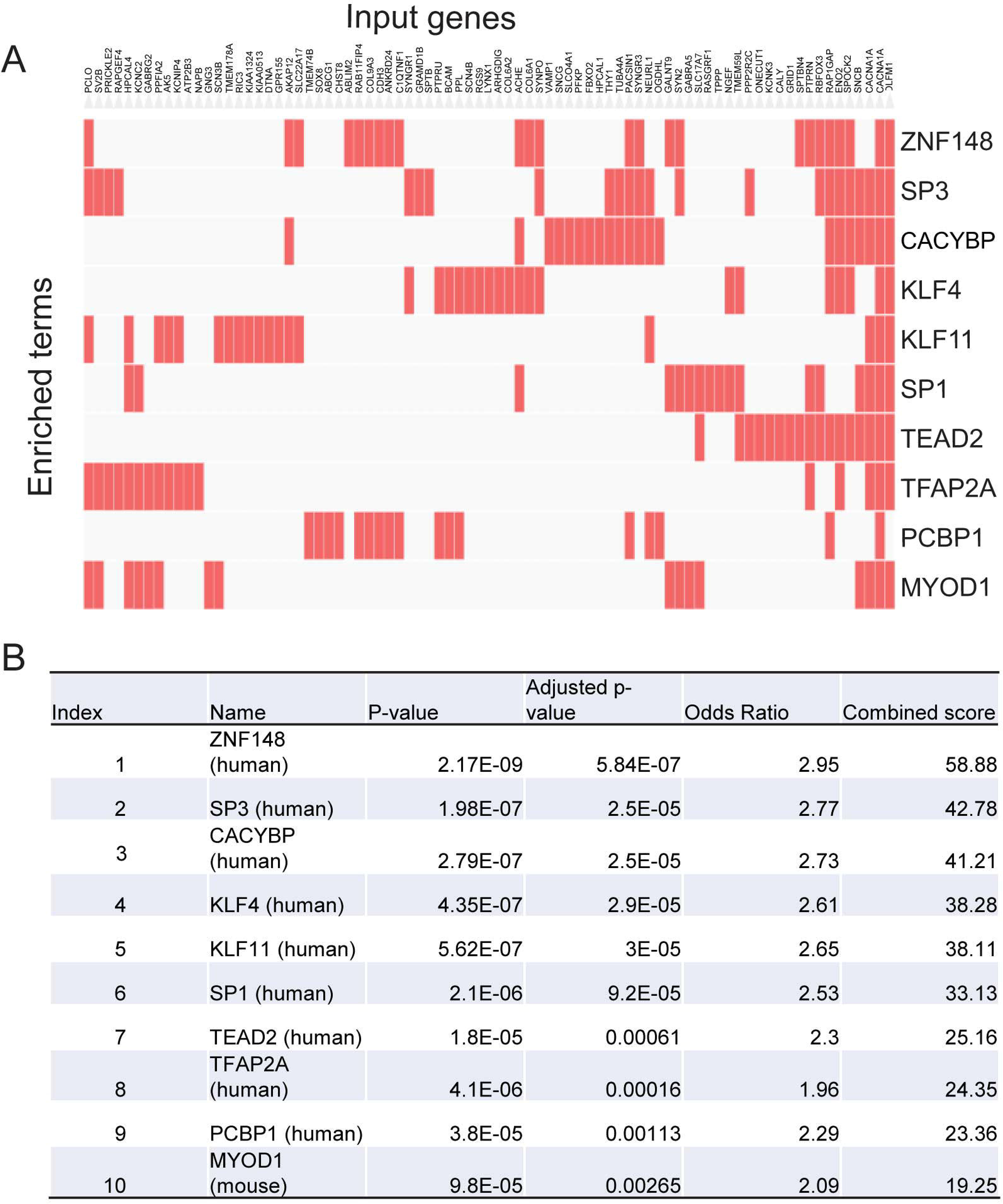
TF binding site enrichment analysis. **(A)** and **(B)** TF binding site enrichment analysis on downregulated genes in *OTUD7A* LoF mutants by Enrichr. The input gene set included all differentially expressed genes that had downregulated expression in *OTUD7A* LoF mutants with *p* < 0.01. Position weight matrices (PWMs) from TRANSFAC and JASPAR collection were used as the binding motifs for the detection at the gene promoters. Index: rank of the TF enriched.

## Supplementary Tables’ Captions

Table S1. All rare putative LoF mutations (stop-gain or frame-shift indels) with sample minor allele frequency (MAF) < 0.005 and their minor allele counts in the sequenced SZ cases and controls.

Table S2. Minor allele counts of rare LoF mutations (sample MAF < 0.005) by gene in SZ cases and controls. To calculate the ratio of allele counts in cases and controls, 0.01 was added to the control counts as denominator. Genes were ranked by their pLI score.

Table S3. SZ cases with rare (MAF < 0.5%) LoF mutations in *OTUD7A* (Refseq NM130901.3) identified by exonic sequencing of CNV genes. Please note that mutation c.1465_1469delAACGG was later verified by Sanger sequencing as a 9-bp in-frame deletion c.1465_1473delAACGGCAAG/p.489-491(AspGlyLys).

Table S4. OTUD7A LoF mutations in SCHEMA (meta_ENSG00000169918_variants_2021_08_18_16_42_15). AC, allele count; AN, total allele numbers; AF, Allele frequency.

Table S5. RNA-seq analysis of DEGs between neurons from wild type and mutant hiPSC lines

Table S6. GSEA using WEB-based Gene SeT AnaLysis Toolkit (WEBGestalt) revealed that the downregulated DEGs in the mutant iNs showed strong enrichment for the genes associated with synapse and synaptic signaling. Top 10 most enriched GO terms were listed for each category. BP, biological process; CComp, cellular components.

Table S7. Sequences of the used guide RNAs (gRNAs) and ssODN for CRISPR SNP editing and PCR primers for mutation confirmation, validating the on-target SNP editing and the absence of off-target editing

